# Cell Type-Specific Regulation by a Heptad of Transcription Factors in Human Hematopoietic Stem and Progenitor Cells

**DOI:** 10.1101/2023.04.18.537282

**Authors:** Shruthi Subramanian, Julie A.I. Thoms, Yizhou Huang, Paola Cornejo, Forrest C. Koch, Sebastien Jacquelin, Sylvie Shen, Emma Song, Swapna Joshi, Chris Brownlee, Petter S. Woll, Diego Chacon Fajardo, Dominik Beck, David J. Curtis, Kenneth Yehson, Vicki Antonenas, Tracey O’ Brien, Annette Trickett, Jason A. Powell, Ian D. Lewis, Stuart M. Pitson, Maher K. Gandhi, Steven W. Lane, Fatemeh Vafaee, Emily S. Wong, Berthold Göttgens, Hamid Alinejad Rokny, Jason W.H Wong, John E. Pimanda

## Abstract

Hematopoietic stem and progenitor cells (HSPCs) rely on a complex interplay of transcription factors (TFs) to regulate differentiation into mature blood cells. A heptad of TFs - FLI1, ERG, GATA2, RUNX1, TAL1, LYL1, LMO2 - bind regulatory elements in bulk CD34+ HSPCs. However, whether specific heptad-TF combinations have distinct roles in regulating hematopoietic differentiation remained unknown. We mapped genome-wide chromatin contacts and TF binding profiles in HSPC subsets (HSC, CMP, GMP, MEP) and found that heptad occupancy and enhancer-promoter interactions varied significantly across cell types and were associated with cell-type-specific gene expression. Distinct regulatory elements were enriched with specific heptad-TF combinations, including stem-cell-specific elements with ERG, and myeloid- and erythroid-specific elements with combinations of FLI1, RUNX1, GATA2, TAL1, LYL1, and LMO2. These findings suggest that specific heptad-TF combinations play critical roles in regulating hematopoietic differentiation and provide a valuable resource for development of targeted therapies to manipulate specific HSPC subsets.

## Introduction

Hematopoietic stem cells (HSCs) maintain production of circulating blood cells via their capacity to either self-renew or differentiate to mature cell types (Doulatov et al., 2012). The most primitive HSCs have multilineage potential but can give rise to progenitor cells with increasing lineage restriction. Although single cell analyses have suggested that differentiation occurs over a continuum rather than in discrete leaps (Corces et al., 2016; Karamitros et al., 2018; Laurenti and Gottgens, 2018; Setty et al., 2019; Velten et al., 2017), relatively pure populations which correspond to intermediate progenitor stages can be prospectively isolated based on cell surface markers (Buenrostro et al., 2018).

Changes in cell identity that occur across differentiation trajectories are directly related to altered transcriptional programs (Buenrostro et al., 2018; Novershtern et al., 2011) which are in turn controlled by lineage specific gene regulatory networks (GRNs) (Bordukalo-Niksic et al., 2022; Xu et al., 2012). At the simplest level, GRNs are comprised of genes, their associated regulatory elements (promoters and *cis*-regulatory elements (CREs) such as enhancers), and transcriptional regulators, including transcription factors (TF), which bind these elements (Davidson, 2010; Thoms et al., 2019). Accessibility of regulatory elements is controlled by various chromatin modifications, and the DNA sequence of such elements at least partially determines the specific TFs that can bind (Lara-Astiaso et al., 2014; Nasrallah et al., 2016; Thompson et al., 2022; Zaret, 2020). A further layer of control is imposed by chromatin organization into topologically associated domains (TADs) (Dixon et al., 2012; Lieberman-Aiden et al., 2009). Interactions between promoters and their CREs, mediated by chromatin loops and complexes of transcriptional regulators, modulate GRNs and therefore cell identity (Dixon et al., 2015; Novo et al., 2018).

We have previously shown that seven TFs (heptad: FLI1, ERG, GATA2, RUNX1, TAL1, LYL1, and LMO2), all of which are known regulators of healthy and leukemic haematopoiesis (Cai et al., 2000; Curtis et al., 2012; Hahn et al., 2011; Li et al., 2015; Mangan and Speck, 2011; Marcucci et al., 2005; Oram et al., 2010; Pimanda et al., 2007; Thoms et al., 2011; Vicente et al., 2012), bind combinatorially in bulk CD34^+^ hematopoietic stem and progenitor cells (HSPCs) (Beck et al., 2013) and in leukemias (Diffner et al., 2013; Mandoli et al., 2014; Mandoli et al., 2016; Sotoca et al., 2016; Thoms et al., 2021). In healthy HSPCs, heptad combinatorial binding occurs at regulatory regions associated with genes involved in stem cell maintenance and function, and also occurs at heptad CREs such that heptad genes form a highly interconnected regulatory circuit (Beck et al., 2013; Wilson et al., 2010). However, the study of GRNs in HSPCs is hindered by cell-type heterogeneity within the CD34^+^ population and the absence of experimental evidence linking promoters to distal regulatory elements.

To address these issues and further our understanding of heptad-centred GRNs in blood development, we sorted CD34^+^ HSPCs into HSC/multipotent progenitor (MPP), common myeloid progenitor (CMP), granulocyte macrophage progenitor (GMP), and megakaryocyte erythroid progenitor (MEP) cell types. We then used chromatin immunoprecipitation followed by sequencing (ChIP-seq) targeting 10 TFs [heptad, PU.1, CTCF, STAG2] and 3 histone modifications [H3K27ac, H3K4me3, H3K27me3], Hi-C, and H3K27ac-HiChIP in each of these cell types to chart the regulatory landscape of human HSPC differentiation. Combinatorial binding of heptad TFs was observed in all sorted populations, although specific patterns of chromatin occupancy differed between cell types. Heptad promoter looping to putative enhancers was variable across cell types, and in many cases combinatorial binding was observed at CREs in immature cells prior to formation of loops in more mature progenitors. Genome-wide occupancy of heptad TFs was also variable across cell types, with distinct sets of CREs enriched for heptad binding in MEP compared to GMP. This variation was at least partially due to sequence motifs in the CREs, with motif composition sufficient to predict cell type with high sensitivity and specificity.

## Results

### Genome wide patterns of heptad factor binding in HSPCs

Primary mobilised human CD34^+^ HSPCs were sorted into HSC/MPP (referred to hereafter as HSC), CMP, GMP, and MEP (Figure 1A, Figure S1A), purity checked with colony assays (Figure S1B), and fixed for downstream assays (Figure 1B, Table S1). The cell populations were chosen to span the differentiation trajectory from early multipotent stem cells (HSC) through to progenitors committed to the myeloid (GMP) or erythroid (MEP) lineages.

**Figure 1.**
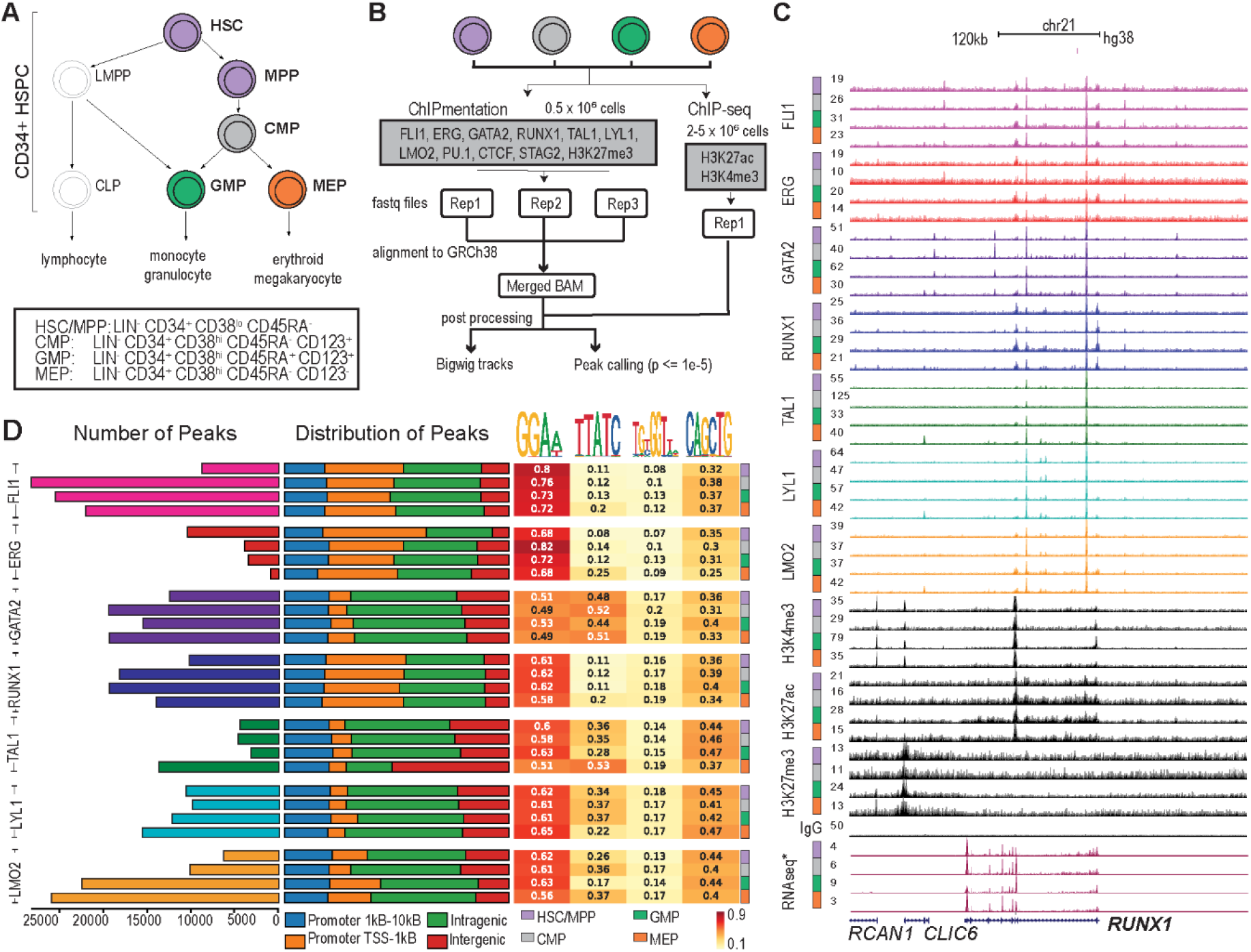
Genome wide patterns of heptad transcription factor binding in fractionated primary human HSPCs. **A)** Human MNCs were isolated from G-CSF-stimulated donors or patients with a non-hematologic malignancy before being enriched for CD34 expression using MACS and further sub-fractionated into individual stem and progenitor cells based on surface marker expression using FACS (colored cells are those studied in this manuscript). **B)** The workflow and analysis pipeline followed for ChIPmentation and ChIP-seq experiments. **C)** UCSC browser track at the *RUNX1* locus (GRCh38 chr21:34,627,969-35,209,177) showing the RPKM-normalized signal from FLI1, ERG, GATA2, RUNX1, TAL1, LYL1, and LMO2, along with H3K4me3, H3K27ac, H3K27me3, IgG (control), and publicly available RNA-seq tracks (GSE75384) for the four cell types. **D)** Characterization of identified peaks. Number of TF peaks were identified by macs2 (p-value ≤ 1e-5) and their overall distribution along the genome (as percentages of total peaks identified) is shown. Each peak was assigned as either promoter-like (proximal [orange] or distal [blue], based on its distance from the TSS), intragenic [green], or intergenic [red], and enrichment calculated within peaks for the known ETS, GATA, RUNX, and E-Box motifs.

High quality ChIP data were obtained for all heptad factors (Figure 1C, Figure 1D, Table S2). The total number of binding peaks was highly variable between heptad factors and across cell types (Figure 1D). We observed cell-type-specific trends consistent with previously described expression patterns and known biology. For example, the highest number of ERG peaks were in HSCs consistent with the role of ERG in maintaining the stem cell state (Knudsen et al., 2015; Tursky et al., 2015), while the highest number of TAL1 peaks were in MEP, consistent with its role in erythroid development (Elwood et al., 1998) (Figure 1D). The distributions of individual TFs across genome features were generally conserved across cell types, but peak distribution differed between heptad factors. For example, FLI1, ERG, and RUNX1 peaks were located at both promoter and non-promoter regions (Figure 1D), while GATA2, TAL1, LYL1, and LMO2 peaks were predominantly located at inter- and intragenic regions. The TAL1 peak distribution in MEP had a unique pattern with an increased proportion of peaks found at intergenic regions. Motif enrichment analysis showed enrichment for ETS (GGAA) and E-Box (CANNTG) motifs in TF occupied regions from all factors. FLI1, ERG, and RUNX1 peaks were highly enriched for the ETS motif, while GATA2, TAL1, LYL1, and LMO2 peaks showed additional enrichment for the GATA motif (GATA), which was particularly prominent for GATA2 and TAL1 in MEP. Overall, we observed conserved patterns of heptad binding, but also distinct differences between factors and across cell types, consistent with dynamic remodeling of the heptad network across the HSPC differentiation trajectory.

### Combinatorial binding of heptad transcription factors is cell-type specific

We have previously described combinatorial binding of heptad factors in bulk human HSPCs (Beck et al., 2013). Here, we extend these observations to specific cell populations. We quantified ChIP peaks containing all possible combinations of two or more heptad factors (Figure 2Ai, ii), and evaluated the probability these occurred by random chance in each cell type (Figure 2Aiii; higher z-score indicates lower probability that the combination occurs by random chance). As observed in bulk HSPCs (Beck et al., 2013), combinatorial binding of all seven factors was the most significant event in any of the four cell types (z-scores for all seven factors; HSC: 8199.94, CMP:6314.92, GMP:3543.93, MEP:1877.48). Pairwise combinations generally had low significance scores, with a few exceptions such as ERG/RUNX in HSCs. In general, higher order complexes had higher significance scores, with five of the seven possible six-factor complexes showing highly significant binding in at least one cell type. Specific combinations which broadly matched known TF function were also observed. For example, five- or six-factor combinations lacking the erythroid regulators GATA2, TAL1, or both, had low significance scores in MEPs (Figure 2A, stars) (Johnson et al., 2020; Thoms et al., 2021; Wontakal et al., 2012). Overall, we find that combinatorial binding of heptad TFs is a general feature of stem and progenitor cells, with some combinations restricted to specific cell types.

**Figure 2.**
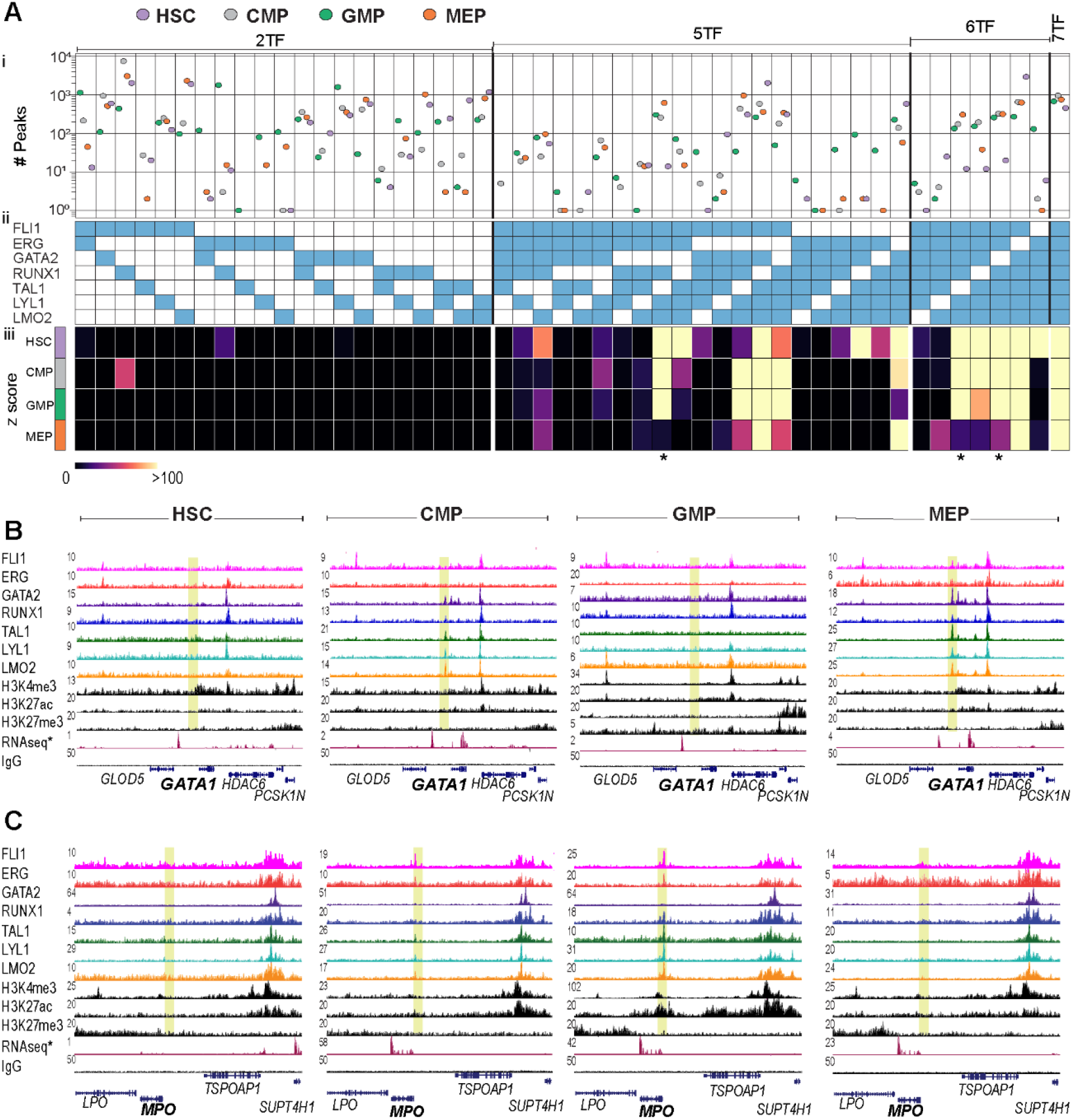
Combinatorial binding of heptad transcription factors is cell-type specific. **A)** A composite graph with three components: i) number of combinatorial binding peaks identified in the four cell types, for ii) combinations of 2, 5, 6, and 7 heptad factors and iii) heatmap showing z-scores for the combinations presented in (ii). Star indicates combinations lacking GATA2 and/or TAL1. **B-C)** UCSC browser tracks showing RPKM-normalized signal tracks of the heptad factors, H3K4me3, H3K27ac, H3K27me3, RNA-seq (public data: GSE75384), and IgG (control) in HSC, CMP, GMP, and MEP (left to right), at **B)** the *GATA1* locus (GRCh38 chrX:48,724,037-48,839,866), a gene vital for erythroid lineage specification, and at **C)** the *MPO* locus (GRCh38 chr17:58,238,087-58,348,896), a gene specific to the monocytic lineage.

Given the cell-type-specific occurrence of some TF combinations we predicted that dynamic formation of TF complexes might play a role in priming CREs for subsequent activation. We explored the binding of heptad factors at promoters of two lineage-specific genes: the erythroid regulator *GATA1* and the monocyte gene *MPO,* neither of which showed heptad binding in HSCs. At the *GATA1* promoter (Figure 2B; yellow region), binding of GATA2, RUNX1, TAL1, LYL1, and LMO2 was observed in CMP. As cells transitioned to MEP, binding peaks became more prominent and now included FLI1 and ERG. However, there was essentially no heptad binding at the *GATA1* promoter in GMP. At the *MPO* promoter (Figure 2C; yellow region), FLI1, ERG, TAL1, LYL1, and LMO2 binding was observed in CMP. Binding peaks became more prominent as cells transitioned to GMP, with RUNX1 also bound, but no TF binding was observed at the *MPO* promoter in MEP. Taken together our data suggests that distinct patterns of heptad TF binding at *cis*-regulatory regions may prime the genomic landscape of human blood stem cells towards either an erythroid or myeloid fate.

### Heptad regulatory circuits are remodelled during myeloid progenitor development

Genes encoding the heptad TFs form a densely interconnected regulatory circuit in bulk HSPCs (Beck et al., 2013), and chromatin accessibility at heptad gene CREs is sufficient to predict cell identity across blood development (Thoms et al., 2021). To broaden our understanding of genome organization and connectivity at heptad loci across HSPC development we performed HiC and H3K27ac HiChIP experiments on the same four sorted cell populations. While more than half of the genome compartments remained stable across the four cell types, some compartments underwent either B to A, or A to B switching during the transition from CMP to lineage committed progenitor (Figure S2Ai). Notably, a series of regions underwent compartment switching in GMP only, with corresponding changes in average H3K27ac signal in these regions (Figure S2Aii, iii). TAD boundaries were highly conserved between cell types (Figure S2B), and HiC contact matrices around the gene loci of the heptad factors were also highly similar (Figure S3).

H3K27ac HiChIP experiments generated thousands of significant interactions with false discovery rate ≤ 0.01 (HSC: 26,210, CMP: 8170, GMP: 43,448, and MEP: 32,773). We focused on looping interactions where at least one interacting region had been annotated as a promoter (P). As this experiment enriched for fragments with the H3K27ac active enhancer mark, we consider looped CREs as putative enhancers (E). We further filtered promoter-enhancer (P – E) loops based on the presence of an ATAC peak at the distal enhancer and integrated these with our ChIP-seq data to create regulatory network maps at each heptad gene locus for each cell type.

Promoter-enhancer loops corresponding to the heptad spanned wide genomic regions; around 500kb for *FLI1*, *GATA2*, and *TAL1* and 1-2Mb for *ERG*, *RUNX1*, *LYL1*, and *LMO2* (Figure 3Ai,ii, Figure S4). Overall, we detected multiple putative enhancers for the heptad genes *ERG* (−610, - 410, -230, +85, +88, +191, and +1200 [+1.2K]), *FLI1* (+27, +32, and +64), *GATA2* (−123, -92 and +4), *RUNX1* (−880, +22, +100, +110, +141, and +161), *TAL1* (−101, −82, −25, +0.5, +14, and +45), *LYL1* (−744, −50, +165, and +310), and *LMO2* (−570, −100, −67, −61, −51, −23, −22, −15, and −12) (Figure 3Ai, ii, Figure S4, Table S3). Although some of these regions have been previously described in humans and/or in mice (Bee et al., 2009; Chan et al., 2007; Gottgens et al., 2010; Gottgens et al., 2002; Landry et al., 2009; Nottingham et al., 2007; Sanchez et al., 1999; Schutte et al., 2016; Sinclair et al., 1999; Wilson et al., 2009; Wozniak et al., 2007), others were novel (Table S3). Two previously described heptad enhancers were not directly looped to the corresponding gene promoter in our data set (*FLI1* - 15, *GATA2* -117) (Beck et al., 2013; Johnson et al., 2015; Moignard et al., 2013), although we did observe looping between *GATA2* -117 and other putative enhancers in MEPs (Figure S4Bi). In addition, we observed that looping at heptad genes generally increased in complexity in GMP and MEP compared to HSC, with the lowest level of looping observed in CMP (Figure 3Ai, Figure S4). This additional complexity often included extensive looping to additional CREs which were not directly connected to heptad promoters, suggesting that heptad gene expression is fine-tuned by highly interconnected cell-specific enhancer communities (eg/ TAL1 locus in MEP, LYL1 locus in GMP and MEP).

**Figure 3.**
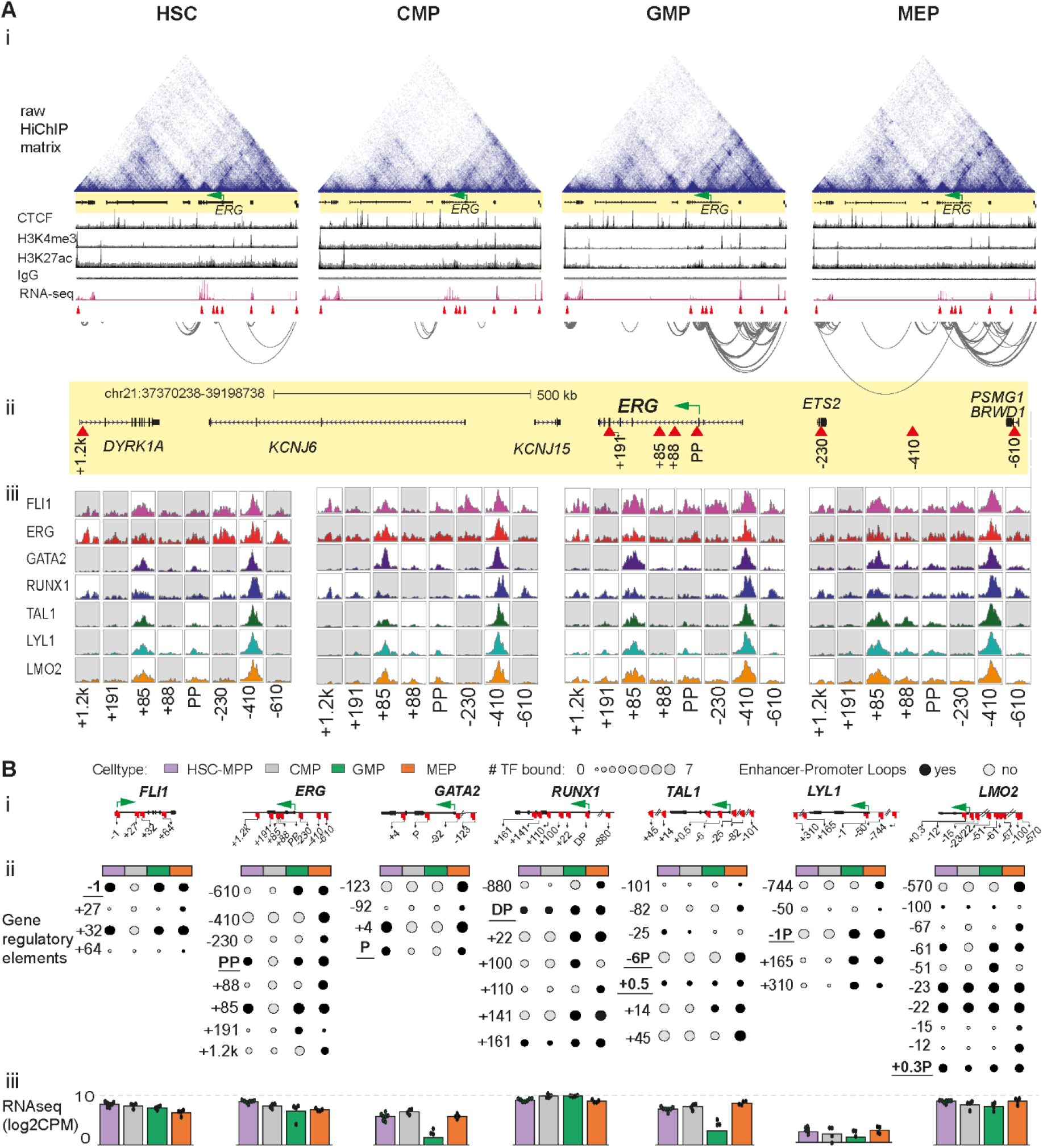
Heptad regulatory circuits are remodelled during myeloid progenitor development. **A)** Step-wise identification of potential regulatory regions interacting with the ERG promoter: i) Raw HiChIP contact matrix, CTCF, H3K4me3, H3K27ac, IgG, RNA-seq, and significant H3K27ac HiChIP interactions (FDR ≤ 0.01) at the *ERG* locus (GRCh38 chr21:37370238-39198738). The *ERG* promoter is indicated by the green arrow (only those HiChIP interactions where both interacting ends were found at the given locus are shown). ii) Magnified view of the *ERG* locus, with regulators identified to loop to the *ERG* proximal promoter shown as red triangles. iii) FLI1, ERG, GATA2, RUNX1, TAL1, LYL1, and LMO2 peaks at the defined regulators in each individual cell type. The peaks shown are RPKM-normalised and white boxes indicate presence of a computationally called ChIP-seq peak at the specific region. **B)** Summary plot of gene regulatory interactions across the heptad genes: i) Individual heptad gene loci with identified regulators indicated by red markers. ii) Dot plots showing regulatory regions as rows and the four cell types as columns, with size of the dot indicating number of heptad factors bound and bold color indicating the presence of an active regulatory link to the promoter (using H3K27ac HiChIP). Promoters are underlined. iii) Bar plots with individual replicates showing average log2 counts of relevant heptad gene expression in the four cell types (GSE75384).

Most of the directly looped elements showed heptad factor binding in at least one cell type (Figure 3Aiii, Figure S4). Some core CREs showed extensive heptad binding in all cell types. This group included known and functionally validated enhancers such as *ERG* +85, *GATA2* +4, *RUNX1* +22, and *TAL1* +45 (Table S3), plus novel distal CREs such as *ERG* -410 and *LMO2* -570 that can now be linked to heptad genes based on our HiChIP data. Integration of HiChIP and ChIP data at heptad gene loci showed diverse patterns of looping and TF binding across the four cell types (Figure 3Bi,ii). At the *ERG* locus, the *ERG +85* enhancer was linked to the proximal promoter in HSC, GMP, and MEP, while other *ERG* elements showed promoter looping in GMP and/or MEP. (Figure 3Bii). Furthermore, heptad TF binding often occurred in HSCs with subsequent promoter looping of that element in more differentiated cell types (e.g./ *GATA2 −123*, *TAL1 +45*, and *LMO2 -570*). However, these epigenetic changes did not directly correspond to the steady state transcriptional output of heptad genes, which was relatively stable across the four cell types (Figure 3Biii).

Observations made in bulk HSPCs indicate that heptad genes regulate themselves and each other via a densely interconnected auto-regulatory circuit (Beck et al., 2013). Using our expanded set of heptad CREs we constructed network connectivity maps for each cell type to visualize dynamic connections over developmental time (Figure S5). These maps highlight the previously unrealized complexity of heptad regulation that underlies steady state gene expression of these key transcriptional regulators. Overall, our data set allowed us to observe extensive remodeling of the regulatory connections within and between individual heptad genes during hematopoiesis and increases our understanding of the complex network regulating heptad genes during hematopoiesis.

### The role of heptad TFs in regulating lineage specific gene expression

We next asked whether cell-type-specific heptad factor chromatin occupancy was associated with cell-type-specific transcriptional output. We identified regions with cell-type-enriched binding of at least 2 heptad TFs (Differentially Enriched for Heptad [DEH]) and used HiChIP data to link these regions to specific genes (Genes associated with DEH [DEHG])(Figure 4A, Table S4A). Approximately half of the regions were promoter-like (up to 10 kb upstream of a TSS); the remainder were considered to be distal regulatory elements. To characterize candidate regulatory elements (REs) and their associated genes we conducted gene set enrichment analysis (GSEA), ingenuity pathway analysis (IPA), and single-cell analysis. Genes linked to DEH regions in HSC/GMP/MEP had greater expression in their respective cell types compared to the other two cell types (Figure 4B). Furthermore, genes differentially bound in GMP were enriched for signaling pathways previously linked to myelopoiesis and granulopoiesis (Figure 4Ci) (Puig-Kroger et al., 2010; Ward et al., 2000a; Ward et al., 2000b) while genes linked to differentially bound regions in MEP were enriched for pathways linked to erythropoiesis (Figure 4Cii) (Dzierzak and Philipsen, 2013; Wendling, 1999). Only 16 HSC-specific genes were identified precluding pathway analysis. However, these genes included known stem cell regulators such as *HOXB1* (Table S4A).

**Figure 4.**
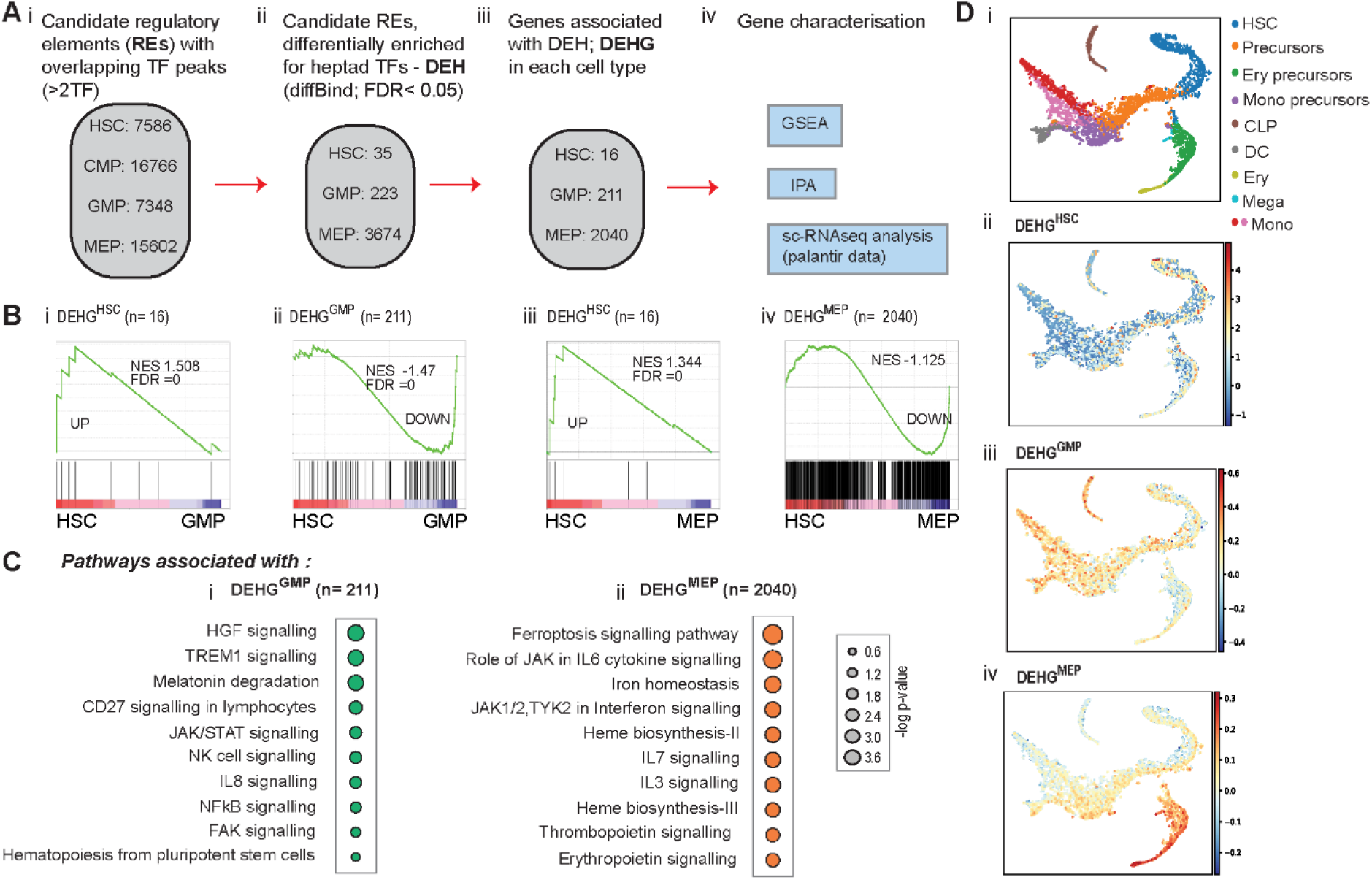
The role of heptad transcription factors in regulating lineage-specific gene expression. **A)** Schematic of the bioinformatic strategy used to derive regions showing differential heptad factor binding: i) Candidate regulatory elements (REs) with binding of at least two heptad factors were chosen in the four cell types; and ii) DiffBind was used to filter for regions showing differential enrichment for heptad factors with an FDR < 0.05. To perform DiffBind analysis only HSC, GMP, and MEP populations were chosen. iii) These DEH (differentially enriched for heptad) regions were linked to genes either directly (present across a 10 kb promoter region) or indirectly (distal links using significant [FDR < 0.01] H3K27ac-HiChIP interactions), and iv) used as input for multiple characterisation assays. **B)** GSEA plots showing enrichment of derived gene sets in pairwise gene expression comparisons: i) DEHG^HSC^ (genes linked to DEH regions in HSC) enriched in HSC with respect to GMP, ii) DEHG^GMP^ enriched in GMP with respect to HSC, iii) DEHG^HSC^ enriched in HSC with respect to MEP, and iv) DEHG^MEP^ enriched in MEP with respect to HSC. **C)** Ingenuity pathway analysis performed in DEHGs: i) DEHG^GMP^, and ii) DEHG^MEP^. **D)** Scoring cell-specific DEHGs along a, i) single-cell expression map reveals localized enrichment of expression: ii) DEHG^HSC^, iii) DEHG^GMP^, and iv) DEHG^MEP^.

We also explored expression of DEHG in single cells across the hematopoietic differentiation trajectory (Figure 4Di) (Setty et al., 2019). As predicted, HSC-associated DEHG (DEHG^HSC^) showed enrichment in cells previously annotated as HSC clusters (Figure 4Dii), while DEHG^GMP^ were found enriched in cells annotated as monocyte clusters (Figure 4Diii) and DEHG^MEP^ in cells annotated as belonging to the erythroid lineage (Figure 4Div). Together these data support our hypothesis that heptad occupancy at regulatory elements can be linked to cell-specific transcriptional output.

### Heptad TF at promoters and distal regulators of genes crucial for myeloid and erythroid development

We next asked whether we could detect specific patterns of heptad occupancy at known lineage specific genes, focusing on genes with roles in mature monocyte and granulocyte maintenance (myeloid), genes linked to erythroid cell development and heme metabolism (erythroid), and genes linked to stem cell function (stem cell) (Table S4B). From these lists we focused on genes whose promoters were looped to a putative enhancer in any of our HiChIP datasets. We identified 40 P-E pairs from the myeloid gene set, 91 P-E pairs from the erythroid gene set, and 81 P-E pairs from the stem cell gene set (Table S4B) and used *k*-means clustering of TF binding signals to compare heptad occupancy patterns at each P-E pair in each cell type (Figure 5A, Figure 5B, Figure S6A respectively). Clustered genes showed varying patterns of TF binding across associated promoter and enhancer regions. Genes in cluster 1 (C1) showed TF enrichment at promoters, genes in cluster 2 (C2) showed TF enrichment at enhancers, and genes in clusters 3 and 4 (C3, C4) had TF enrichment at both promoters and enhancers (Figure 5A, Figure 5B, Figure S6A). Furthermore, several TF-specific observations were evident. First, ERG occupancy at stem cell, myeloid, and erythroid genes was generally highest in HSCs, and to a lesser extent GMPs, which is consistent with its known role in maintaining the stem cell state (Figure S6B, full statistics in Table S5) (Knudsen et al., 2015; Thoms et al., 2021; Tursky et al., 2015). Second, FLI1 and RUNX1 were bound across both myeloid- and erythroid-gene associated regions across all differentiation stages (Figure S6C). Third, GATA2, LYL1, and LMO2 have increased occupancy at myeloid- and erythroid-specific regulatory regions during lineage commitment (Figure S6D, full statistics in Table S5). Fourth, high TAL1 occupancy was observed in MEPs and their precursor CMPs, and this occurred at regions linked to erythroid genes and regions linked to myeloid genes (Figure S6E, full statistics in Table S5). Binding in CMPs may indicate a role for TAL1 in priming regulatory regions and recruiting activators or repressors in downstream cell types. Finally, there was a general pattern of heptad TF occupancy at promoters and enhancers of lineage specific genes even in the earliest stem cells, suggesting that heptad factors bind lineage specific regulatory regions prior to lineage commitment and subsequent differentiation.

**Figure 5.**
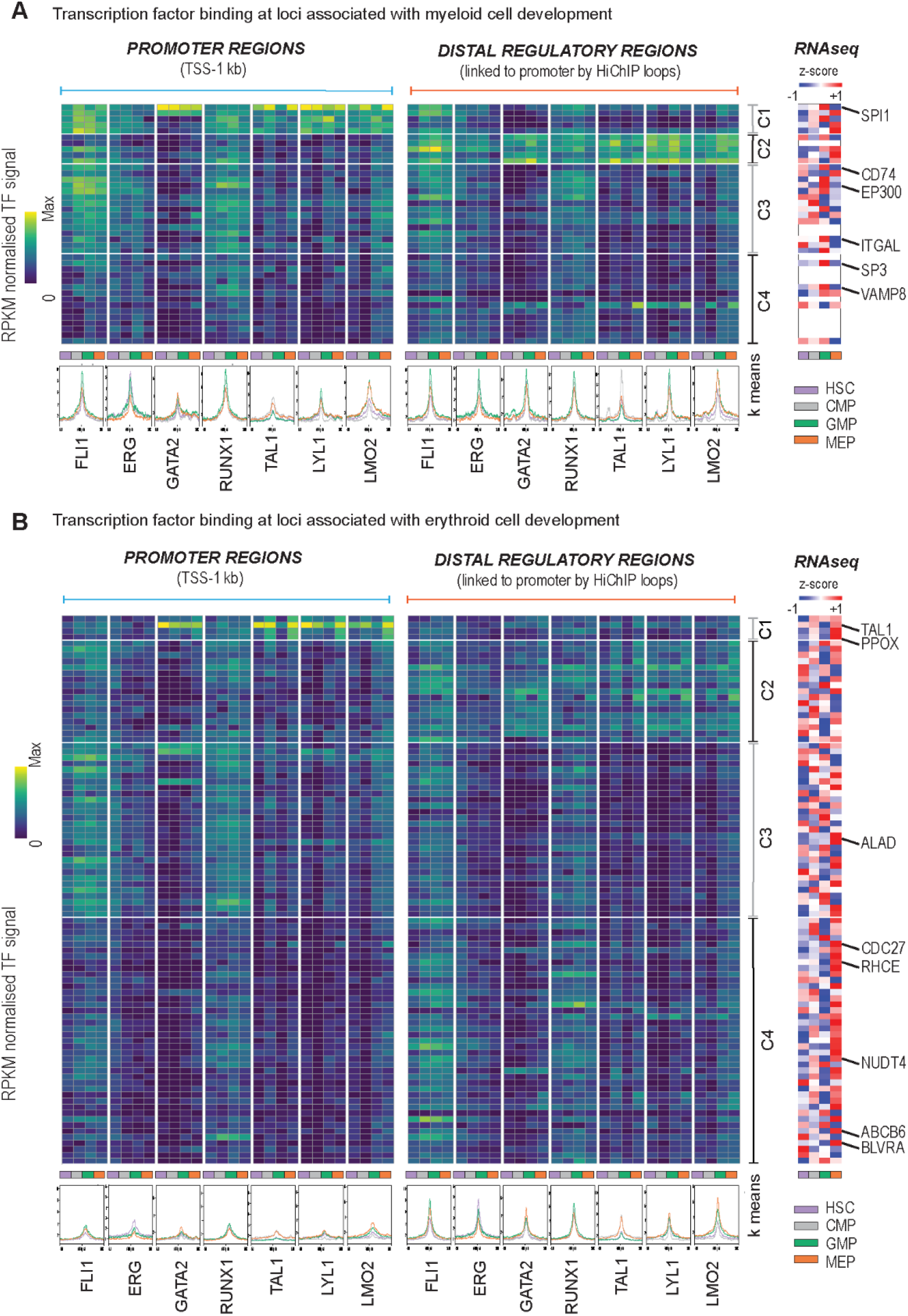
Heptad transcription factors at promoters and distal regulators of genes crucial for myeloid and erythroid cell development. **A)** Genes associated with myeloid development. Left: k-means clustered heatmaps of TF binding intensity at promoters and distal regulatory regions. Profile plots show normalised signal for each TF in each cell type at the regions depicted in the heatmap. Right: z-score normalised heatmaps of RNA-seq counts (GSE75384) for the corresponding gene in each cell type. White rows are genes with no expression values in the dataset. **B)** Genes associated with erythroid development. Left: k-means clustered heatmaps of TF binding intensity at promoters and distal regulatory regions. Profile plots show normalised signal for each TF in each cell type at the regions depicted in the heatmap. Right: z-score normalised heatmaps of RNA-seq counts (GSE75384) for the corresponding gene in each cell type.

### Regulatory regions with cell-type-specific heptad occupancy have distinct epigenetic features

To better understand the underlying mechanisms that regulate dynamic heptad occupancy during blood formation we used our ChIP datasets (heptad, PU.1, CTCF, H3K2lac, H3K4me3, H3K27me3) to annotate and cluster ∼85000 regions previously shown to be accessible in any of our four cell types (Corces et al., 2016). The resulting UMAP projection of the accessibility landscape could be segmented into 13 clusters (Figure 6A). Clusters 1-3 had characteristics of promoters, including high H3K4me3 signal (Figure 6B). Using the associated ChIP signals at each region we could further classify cluster 1 as active promoters (enrichment of H3K27ac and H3K4me3), cluster 2 as bivalent promoters (enrichment of both H3K4me3 and H3K27me3 signal), and cluster 3 as active promoters which were also bound by CTCF. These classifications aligned with ChromHMM annotation of the same regions (Figure 6C) (Ernst and Kellis, 2012). Cluster 4 was enriched for CTCF alone (Figure 6B) and may contain regions primarily involved in 3D genome organization. Clusters 5-11 could be broadly characterized as non-promoter regulatory regions which again aligned with their ChromHMM annotations (Figure 6C). Most of the variations in these regions mapped to variable TF occupancy in specific cell types (Figure S7A), which was particularly pronounced in clusters 8-11 which also showed cell type specific changes in chromatin accessibility (Figure 6B). For example, cluster 9 and 10 both showed high accessibility in MEP compared to GMP, accompanied by high TAL1 and LYL1 occupancy (Figure S7A), suggesting that regulatory regions in these clusters may function in the erythroid lineage.

**Figure 6.**
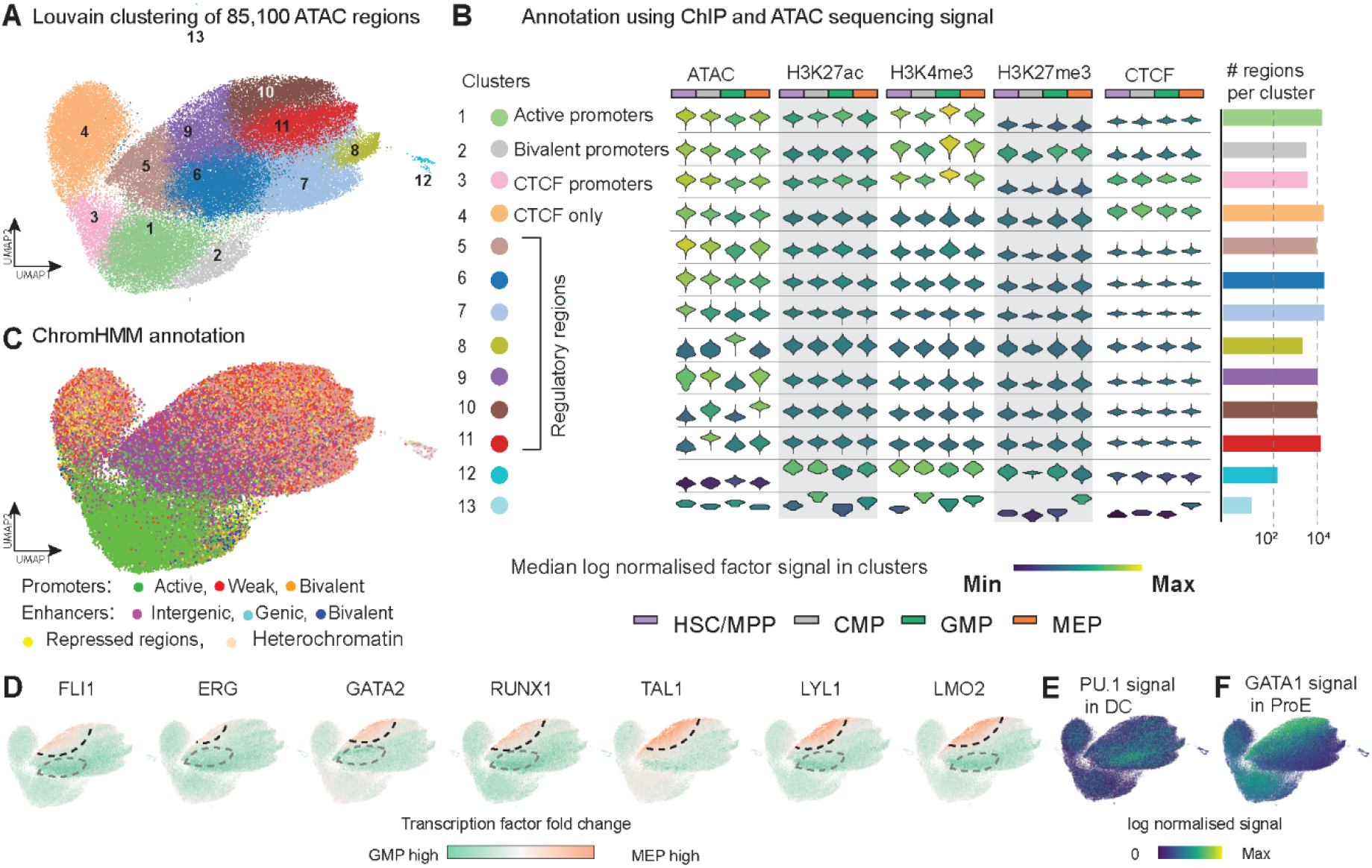
Regulatory regions with cell-type-specific heptad occupancy have distinct epigenetic features. **A)** A UMAP depicting the result of clustering 85,100 accessible regions in HSPCs annotated with ChIPmentation/ChIP-seq signal strengths using the Louvain algorithm. **B)** Individual violin plots of log normalised signal derived from ATAC, 3 histone marks (H3K27ac, H3K4me3, and H3K27me3), and CTCF – accompanied by a bar plot showing the number of regions in each cluster. Inter-cluster signal variability allows annotation of individual clusters based on their regulatory potential. **C)** UMAPs overlaid with ChromHMM annotation of 85,100 individual regions show striking similarity with **B**. **D)** UMAPs coloured based on log2 fold change of binding of the heptad transcription factors in pairwise comparisons between GMP and MEP. MEP- and GMP-specific enrichment of TF binding is identified, and borders demarcated by dashed lines: black (enriched in MEP) or grey (enriched in GMP). **E)** Signal of PU.1 in dendritic cells (DC) (GSE58864) across the clustered regions. PU.1 signal enrichment in dendritic cells mirrors heptad factor enrichment patterns in GMP. **F)** Signal of GATA1 in proerythroblasts (ProE) (GSE36985) across the clustered regions. GATA1 signal enrichment in proerythroblasts mirrors heptad factor enrichment patterns at these regions in MEP.

Visualization of the TF signal across the accessibility landscape revealed variable occupancy patterns across cell types (Figure S7B). FLI1, RUNX1, and ERG had similar distributions across all cell types, with FLI1 and RUNX1 occupancy highest in CMP and GMP, while overall ERG occupancy was reduced in MEP. LYL1 and LMO2 had similar distributions and were enriched in clusters 5/6/7 in GMP and in clusters 9/10 in MEP (Figure S7B). TAL1 was also enriched in clusters 9/10 in MEP with lower binding in all other cell types, while PU.1 was enriched in cluster 5/6/7 in CMP and GMP (Figure S7B). GATA2 had a unique occupancy pattern with enrichment in cluster 5/6 in HSC and GMP and additional occupancy in cluster 9 in CMP and MEP (Figure S7B). This pattern may reflect distinct roles of GATA2 in early hematopoiesis and subsequent erythroid specification (Bresnick et al., 2010).

We then performed pairwise comparisons of TF binding signals in GMP vs MEP, HSC vs GMP, and HSC vs MEP (Figure 6D, Figure S7C). Comparing GMP (Figure 6D, green) to MEP (Figure 6D, orange), there were distinct zones of TF enrichment in GMP (Figure 6D, grey dotted line) and MEP (Figure 6D, black dotted line) for all TFs in both cell types except for TAL1 which showed no region of enrichment in GMP. To further confirm that the regions enriched for heptad TF binding in GMP and MEP represent lineage-specific regulators we mapped the ChIP seq signal from lineage defining TFs in two mature cell types (Figure 6E, F). Regions with high heptad occupancy in GMP showed similar high occupancy of PU.1 in dendritic cells (Figure 6E) (Seguin-Estevez et al., 2014), while regions with high heptad occupancy in MEP showed similar high occupancy of GATA1 in proerythroblasts (Figure 6F)(Xu et al., 2012). Taken together these data show that regulatory regions with heptad occupancy in progenitor populations are regions occupied by lineage specific TFs in more mature cells.

### Cell-type specificity of regulatory elements is encoded in the underlying motif composition

TFs bind DNA via their consensus binding motifs and the sequence and relative locations of motifs in each regulator determine which TF complexes can potentially bind (Inukai et al., 2017). To better understand the enhancer features which underpin lineage specific TF occupancy we selected regions with differential accessibility and TF occupancy in HSC (Figure 7Ai: 3992 regions shown in purple), GMP (Figure 7Bi: 4395 regions shown in green) and MEP (Figure 7Ci: 3469 regions shown in orange) and developed machine learning models to predict cell-type associations based on DNA sequence motifs. All models showed high sensitivity and specificity to associate regions with cell types (Figure 7Aii, Bii, Cii). To understand how specific motifs contribute to assigning regulatory elements to cell-types, we calculated SHapley Additive exPlanations (SHAP) values for each cell type (Figure 7Aiii, Biii, Ciii, Table S6). Distinct combinations of motifs were found to contribute to each cell-type model, and many of these motifs fit the expected profile for cell-specific regulatory regions. For example, ETS motifs had positive SHAP values in the GMP model but negative values in the MEP mode, while GATA motifs had high SHAP values in the HSC and MEP models but negative value in the GMP model. This is consistent with known roles for ETS factors such as PU.1 in driving myeloid differentiation and GATA factors such as GATA1 driving the erythroid lineage (Rosenbauer et al., 2004; Zhang et al., 2000). However, motifs that do not correspond to heptad factors also had high SHAP scores in all cell-type models, reinforcing that heptad factors bind at lineage specific enhancers in the context of larger regulatory protein complexes (Wadman et al., 1997; Wozniak et al., 2008).

**Figure 7.**
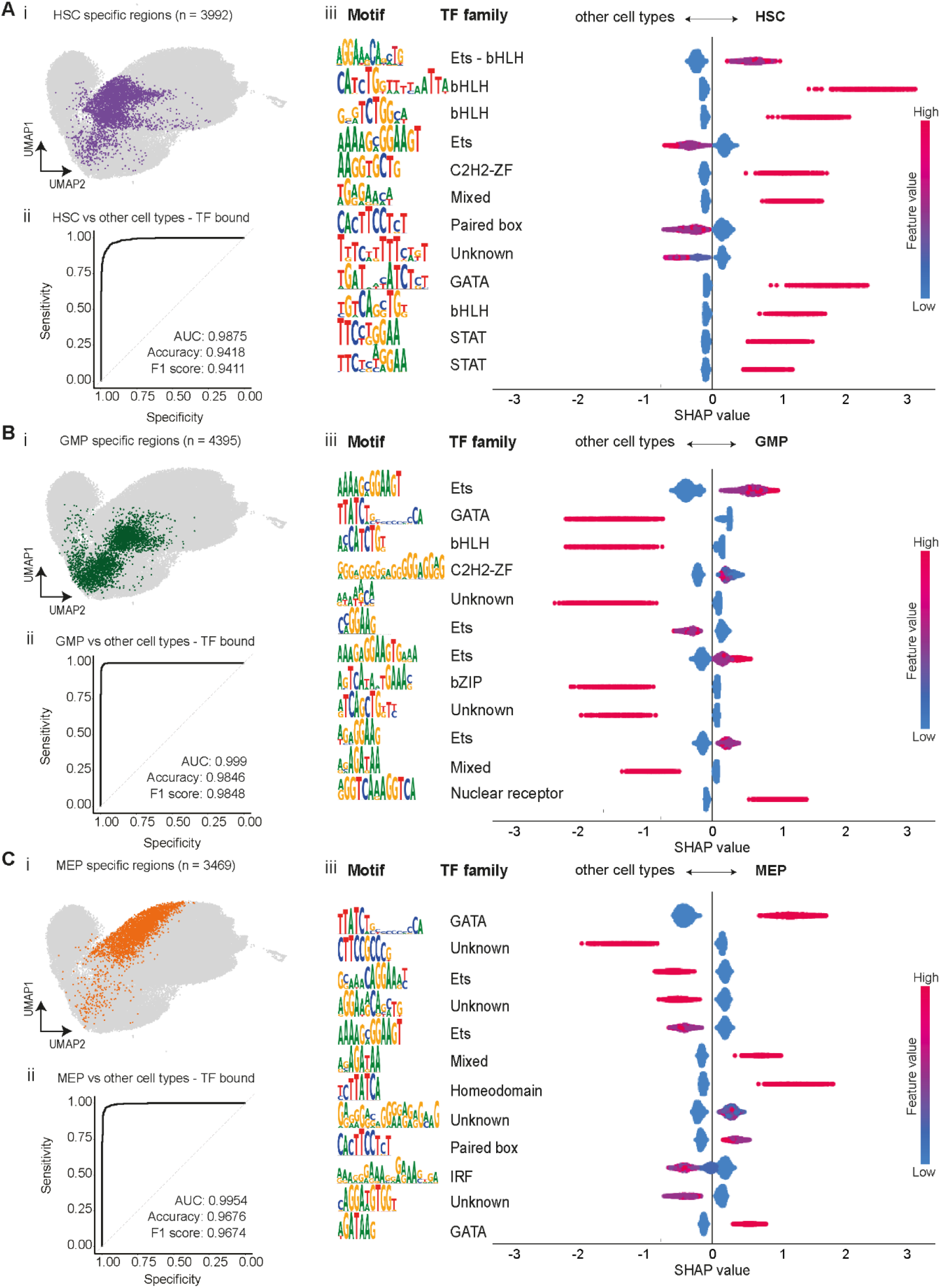
Cell-type specificity of regulatory elements is encoded in the underlying motif composition. **A)** (i) UMAP representation of ATAC-seq regions in CD34^+^ cells (grey) with heptad TF bound HSC specific regions colored in purple. (ii) An XGBoost machine learning model was trained and tested with motif counts from a mixture of regions specified in (i) and background regions, to predict cell type with high accuracy. The ROC curve shows the predictive performance of the constructed model to predict HSC specific regions. (iii) A beeswarm plot depicting the top 12 representative motifs in HSC specific regions - ranked based on their absolute importance in contributing to the predictive model. Each row shows the motif (and canonical TF family if known), and the corresponding SHAP (**SH**aply **A**dditive ex**P**lanations) values for the cell type in question (right) and the others (left). The feature count indicates the normalised motif counts with a range of 0-1. **B)** (i) UMAP representation of ATAC-seq regions in CD34^+^ cells (grey) with heptad TF bound GMP specific regions colored in green. (ii) ROC curve showing the performance of the model to predict GMP specific regions. (iii) A beeswarm plot depicting the top 12 representative motifs in GMP specific regions - ranked based on their absolute importance in contributing to the predictive model. **C)** (i) UMAP representation of ATAC-seq regions in CD34^+^ cells (grey) with heptad TF bound MEP specific regions colored in orange. (ii) ROC curve showing the performance of the model to predict MEP specific regions. (iii) A beeswarm plot depicting the top 12 representative motifs in MEP specific regions - ranked based on their absolute importance in contributing to the predictive model.

## Discussion

In this study we explored genome-wide dynamics of chromatin occupancy and structure in four cell types along the HSC to myeloid/erythroid differentiation axis. Our study revealed that combinatorial binding of heptad TFs was highly dynamic across HSPC subsets and that specific TF combinations were associated with particular cell types. Our study also revealed additional complexity in the interconnected heptad regulatory structure with identification of novel putative enhancers with highly dynamic looping to heptad promotors across the differentiation trajectory. Furthermore, we found additional evidence for the role of heptad TFs in driving cell fate decisions during hematopoiesis, with heptad occupancy defining sets of CREs with cell-type-specific activity.

### Looping complexity, enhancer cross talk, and cell-type-specificity

Analysis of putative CREs that were directly looped to heptad promoter regions revealed a much greater level of regulatory complexity than previously appreciated. Previous characterisation of heptad gene regulation has identified a set of nine key regions that have combinatorial heptad binding in human HSPCs and whose relative accessibility is sufficient to predict cell identity (Thoms et al., 2021). In this study we identified more than 30 putative CREs that were directly looped to heptad promoters in at least one cell type (Figure 3B). While most of the previously known heptad regulatory elements have been tested in functional assays (Table S3), further experimental validation will be required to precisely understand the individual and cooperative roles of specific CREs for each heptad gene (Canver et al., 2015; Xu et al., 2022). Surprisingly, a subset of previously identified enhancers were not looped directly to heptad promoters in this analysis. For example we found *GATA2* −117, an enhancer that is dysregulated in inv(3) AML (Groschel et al., 2014; Yamazaki et al., 2014), was only indirectly looped to the *GATA2* promoter, and was therefore excluded from our network model (Figure S4B, Figure S5, Table S7). However, lack of direct binding does not preclude a role for this enhancer in regulating GATA2. Indeed previous studies have found that enhancer – enhancer interactions can stabilise and amplify TF binding, and that the resulting enhancer communities can drive and coordinate cell-type specific gene expression (Madsen et al., 2020; Robles-Rebollo et al., 2022; Zuin et al., 2022). Consistent with this model, we observed a trend of increasingly complex chromatin looping interactions involving heptad promoters and putative enhancers as cells committed to either erythroid or myeloid lineages (Figure 3B).

Motif-based analysis showed that heptad-occupied CREs contain intrinsic information encoding cell-type-specificity. Among the motifs that contributed to cell-type specificity, many corresponded to heptad factors, but we also observed motifs for additional factors including STAT, interferon-regulatory factor (IRF), and homeodomain TFs (Figure 7, Table S6). While our focus was on heptad factors, it is clear that additional regulators co-occupy CREs and contribute to enhancer specificity (Spitz and Furlong, 2012). The precise molecular mechanisms governing enhancer specificity are still unclear. Recent genome wide screens have revealed that groups of enhancers have specific cofactor dependencies (Neumayr et al., 2022). Promoters also have cofactor dependencies (Haberle et al., 2019), and show differential responsiveness to specific enhancers (Arnold et al., 2017). To date these studies have focussed on transcriptional cofactors with broad function such as P53 and MED14 (Neumayr et al., 2022), and further work is required to understand how heptad factors, general cofactors, and intrinsic features of enhancer and promoters cooperate to regulate transcriptional programs throughout haematopoiesis.

### Therapeutic implications for bone marrow failure syndromes

Numerous bone marrow failure syndromes have an underlying germline genetic cause (Dokal et al., 2022) and may therefore be amenable to gene therapy approaches. Such therapies would ideally target the most primitive HSCs to enable long term clinical benefits. However, ectopic gene expression, particularly in stem cells, may lead to lineage skewing and other unwanted outcomes. Cell-type-specific enhancers are conserved across evolution and can potentially be used to limit gene expression to particular cell populations (Cornejo-Paramo et al., 2022; Goode et al., 2016; Xu et al., 2012). Indeed a recent report has described the use of erythroid-specific *GATA1* enhancers to drive restricted expression of GATA1 to rescue erythroid differentiation in Diamond-Blackfan Anaemia, an approach that is agnostic to the precise genetic mutation in individual patients (Voit et al., 2022). However, identifying enhancers with the appropriate spatiotemporal activity for use in gene therapy vectors is not trivial. While various experimental techniques have been developed to allow massively parallel interrogation of enhancer activity and specificity (Edginton-White et al., 2023; Hrvatin et al., 2019; Xu et al., 2022), our data provides a catalogue of putative enhancers already annotated with respect to spatiotemporal activity that could fast track functional studies.

### Implications for understanding GRN perturbation in AML

GRNs are significantly perturbed in multiple AML subtypes (Assi et al., 2019; Thoms et al., 2019). These network rewiring events can often be attributed to specific translocation events, for example RUNX1-RUNX1T1 (which leads to altered RUNX1 function and chromatin occupancy) (Grinev et al., 2021; Loke et al., 2017; Ptasinska et al., 2012; Tonks et al., 2007), or inv(3) (which leads to dysregulation of GATA2 and EVI1) (Groschel et al., 2014; Yamazaki et al., 2014). Recent high depth sequencing studies have led to an increased understanding of the frequency and significance of small structural variants (SVs) which can mediate enhancer hijacking and subsequent oncogenic transformation (Botten et al., 2022; Montefiori et al., 2021; Ottema et al., 2021; Smeenk et al., 2021; Xu et al., 2022). A key difficulty in interpreting and translating such data is the need to distinguish benign SVs from those which play a role in driving leukemic maintenance, and understanding which genes they directly dysregulate. Our study has characterised thousands of potential CREs across the haematopoietic trajectory, and more importantly has annotated their spatiotemporal connectivity and activity.

### Scope and limitations of the study

The datasets generated in this study – including ChIP-seq targeting 10 TFs [heptad, PU.1, CTCF, STAG2] and 3 histone modifications [H3K27ac, H3K4me3, H3K27me3], Hi-C, and H3K27ac-HiChIP in each cell type – are an invaluable resource for study of healthy and malignant haematopoiesis in humans. While extensive, our dataset is limited to four cell types covering the myeloid, but not lymphoid arm of differentiation. Furthermore, even with the low cell input techniques used in this study, our datasets represent averaged binding and looping probabilities captured across a continuously shifting epigenetic landscape, rather than truly capturing GRNs, which operate at the level of single cells (Davidson, 2010). Our datasets also rely on formaldehyde crosslinking which is limited to amines (Hoffman et al., 2015) and unable to capture very brief interactions that may nonetheless be biologically significant (Schmiedeberg et al., 2009); thus care must be taken when interpreting absence of a signal. Since healthy HSPCs from human donors are in limited supply, technical advances in measuring chromatin occupancy in single cells and small input adaptations of higher resolution HiC-based techniques will be required to improve our ability to fully resolve GRN dynamics during haematopoiesis (Aljahani et al., 2022). Nonetheless, our dataset represents the most extensive epigenetic characterisation of primary human blood progenitors to date and provides important insights for future studies in healthy and leukemic cells.

## Conclusions

Overall, we identified regulatory elements with differential heptad binding over the course of HSPC differentiation and found that occupancy of such regions is associated with cell-type-specific gene expression. Furthermore, heptad occupancy in progenitors may prime specific CREs for subsequent binding of lineage defining TFs such as PU.1 and GATA1. Finally, we find that enhancers with cell-type-specific heptad occupancy share a common grammar with respect to TF binding motifs, and thus combinatorial binding of specific TF complexes is at least partially regulated by features encoded in specific DNA sequence motifs. This data set provides a rich resource to guide future studies of both healthy and leukemic haematopoiesis.

## Acknowledgements

The authors acknowledge members of the Pimanda group for assistance in thawing apheresis bags. The authors thank the South Australian Cancer Research Biobank (SACRB), and all patients who donated samples. For the purpose of open access, the author has applied a CC BY public copyright licence to any Author Accepted Manuscript version arising from this submission. JAIT was supported by the Anthony Rothe Memorial Trust. BG acknowledges funding from Wellcome (206328/Z/17/Z). JEP was supported by grants from the National Health and Medical Research Council of Australia (GNT1139787, GNT2011627, MRF1200271), a translational program grant from the Leukemia Lymphoma Society (LLS)-Snowdome Foundation-Leukaemia Foundation (6620-21), and the Anthony Rothe Memorial Trust.

## Author contributions

Conceptualization: SS, JAIT, JWHW, JEP

Methodology: SS, JAIT, YH, SJ, SSh, ES, CB, PSW, DCF, DB, AT, SWL

Investigation: SS, JAIT, YH, SSh, ES, SJ, DJC, KY, VA, TOB, JAP, IL, SMP, MKG

Formal analysis: SS, JAIT, PC, FCK, FV, EW, BG, HAR, JWHW, JEP

Supervision: JAIT, JWHW, JEP

Writing - original draft: SS, JAIT, JEP

Writing – review and editing: JAIT, JEP

Funding acquisition: JEP

## Key resource table

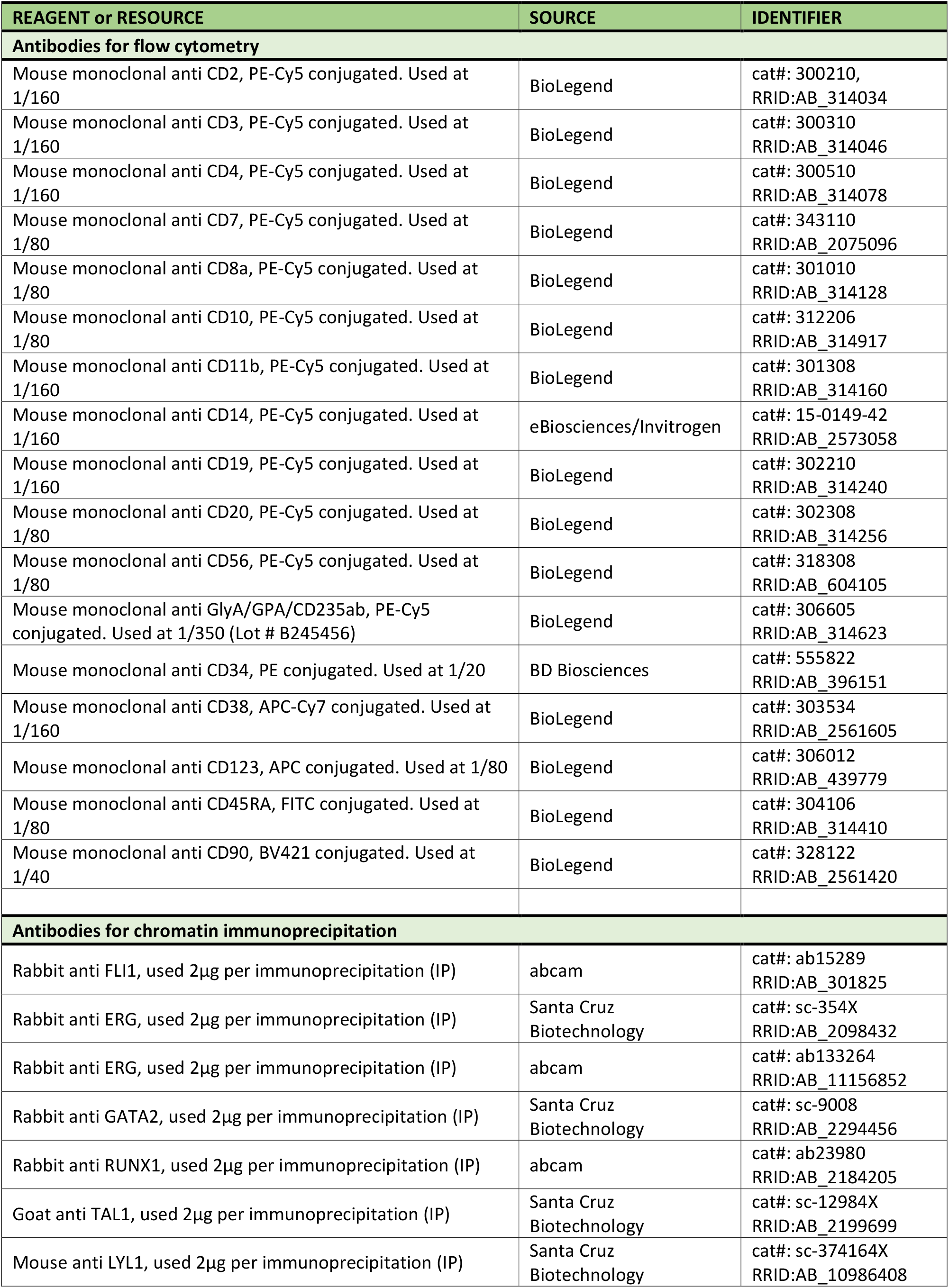
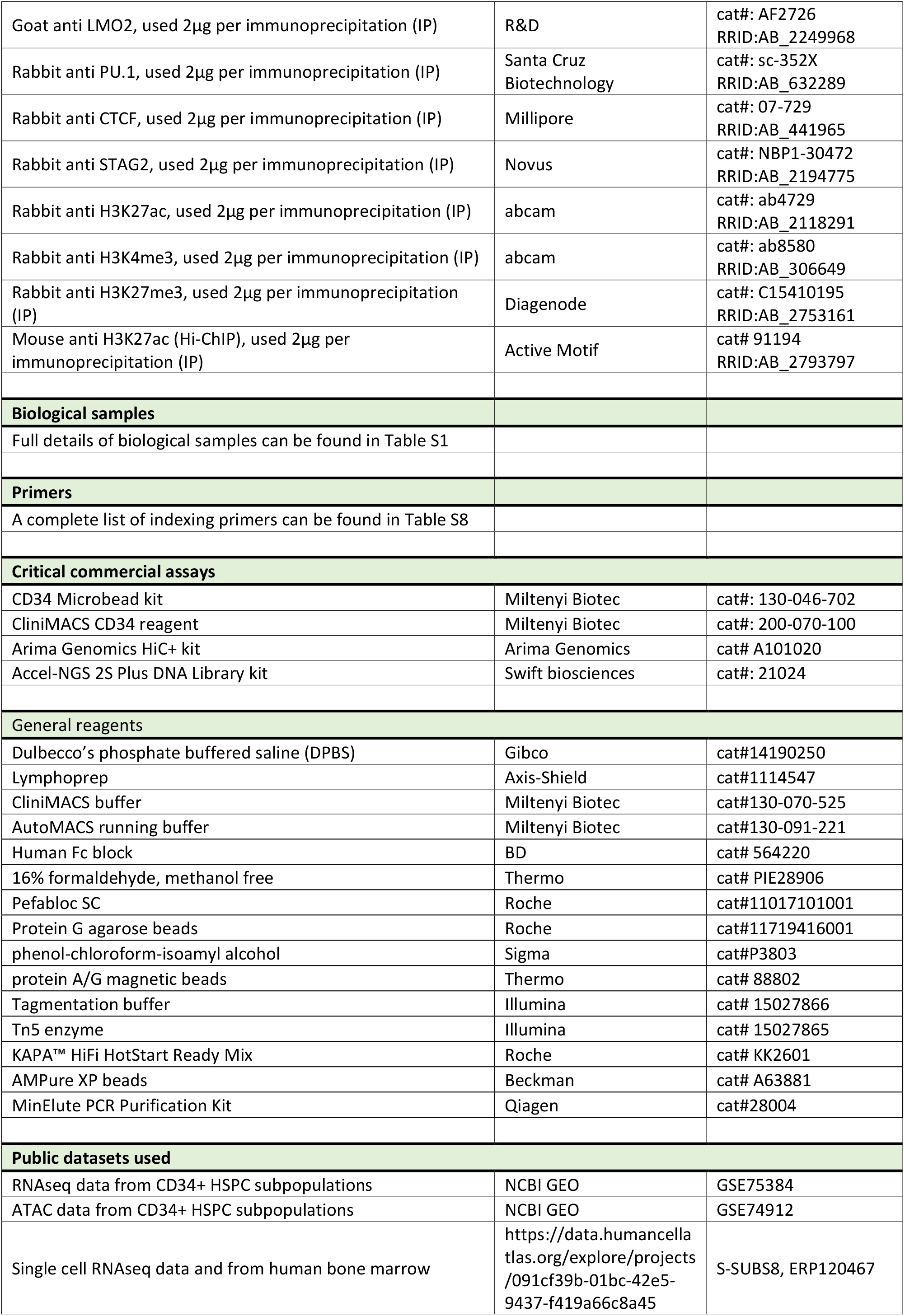
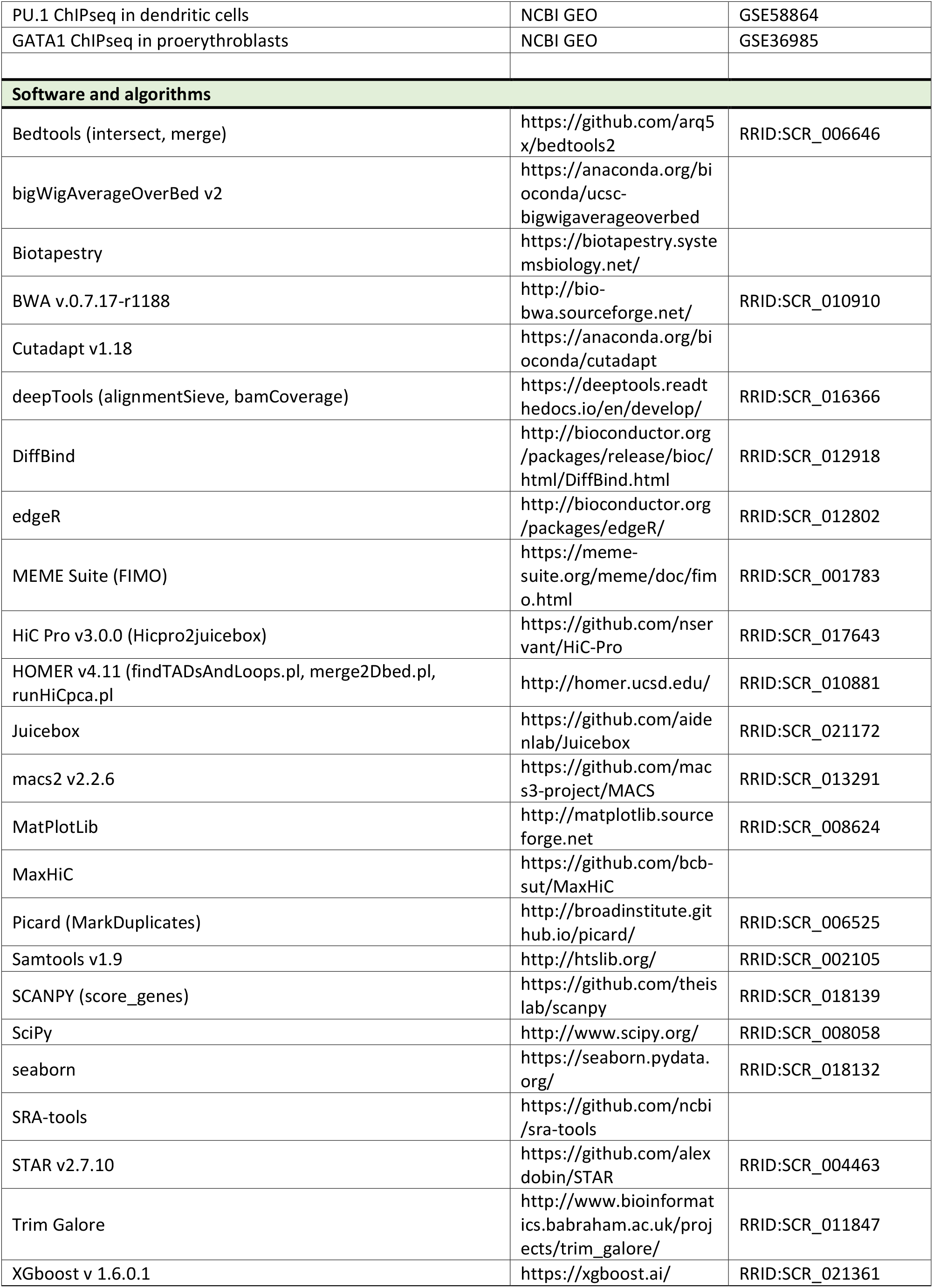

## Methods

### Biological samples

Mobilized peripheral blood samples were collected with patient consent in accordance with the Declaration of Helsinki, and used with institutional ethics approval ref:08/190 from South Eastern Sydney Local Health District, NSW, Australia.

### Isolation of CD34^+^ cells from cryopreserved apheresis packs

Cryopreserved cells were thawed, diluted 1:4 in 2.5% dextran/12.5% human albumin in 0.9% saline, centrifuged 200g/15 min/room temperature (RT) and resuspended in phosphate-buffered saline (DPBS) containing 10% fetal bovine serum (FBS). Cells were underlaid with lymphoprep, centrifuged 800g/30 min/RT, and mononuclear cells (MNCs) collected from the interface and washed with DPBS. MNCs were resuspended in ice cold CliniMACS buffer supplemented with 0.5% human albumin or AutoMACS running buffer then labelled with anti-CD34 microbeads according to manufacturer’s instructions (Miltenyi Biotec). CD34^+^ cells were enriched using either a CliniMACS Plus (Miltenyi Biotec) using standard clinical parameters or an AutoMACS (Miltenyi Biotec) using the program posseld2.

### Labelling and sorting of CD34^+^ cells

CD34^+^ cells were resuspended in FACS buffer (5% FBS/1mM EDTA in DPBS) containing 1/10 diluted FcBlock at a concentration of 10^7^ cells/100 µL and stained on ice for 30 min with a cocktail of antibodies (CD34 subset markers – CD38, CD38, CD123, CD45RA, CD90: Lineage markers (LIN) – CD2, CD3, CD4, CD7, CD8a, CD10, CD11b, CD14, CD19, CD20, CD56, GlyA/GPA/CD235ab). Cells were sorted using a BD FACS ARIA II into the following populations: HSC (includes MPP; LIN^-^, CD34^+^, CD38^lo^, CD45RA^-^), CMP (LIN^-^, CD34^+^, CD38^+^, CD45RA^-^, CD123^+^), GMP (LIN^-^, CD34^+^, CD38^+^, CD45RA^+^, CD123^+^), MEP (LIN^-^, CD34^+^, CD38^+^, CD45RA^-^, CD123^-^) (Figure S1A). Population gates were set using fluorescence minus one controls, and phenotypic purity checks were performed on collected cell fractions.

Functional validation of cell purity was carried out for a subset of experiments. Sorted cells were resuspended 1% methylcellulose supplemented with cytokines as described (Schuller et al., 2007), then plated in triplicate at 500 cells per dish and incubated in a humidified 37°C incubator with 5% CO2 for 14 days. Three major types of colonies were counted: erythroid-lineage (BFU-E) colonies, myeloid-lineage (GM) colonies, and colonies with mixed-potential (GEMM) (Figure S1B).

### Crosslinking and preparation of nuclei

Sorted cells were incubated in freshly prepared 1% formaldehyde in DPBS for 10 min at RT. Crosslinking was quenched by adding glycine to a final concentration of 0.125 M and incubating for 5 min at RT. Subsequent steps were performed at 4°C with cold buffers. Cells were washed then resuspended in cell lysis buffer (10mM Tris-Cl pH 8.0, 10mM NaCl, 0.2% Tergitol, supplemented with 1µg/mL leupeptin, 1mM Pefabloc SC, 10mM sodium butyrate) and incubated on ice for 10 minutes. Nuclei were centrifuged 1450g/10 min/4°C then snap frozen and stored at -80°C for later use.

### Chromatin Immunoprecipitation (ChIP)

ChIP for H3K27ac and H3K4me3 was carried out essentially as described (Diffner et al., 2013). Nuclei (2–5 × 10^6^/IP) were resuspended in 0.65mL nuclei lysis buffer (50mM Tris-Cl pH 8.0, 10mM EDTA, 1% SDS, protease inhibitors), incubated on ice for 10 minutes, with 0.4mL IP dilution buffer (20mM Tris-Cl pH 8.0, 2mM EDTA, 150mM NaCl, 1% Triton X-100, 0.01% SDS) and sonicated for 10 cycles in a Bioruptor Pico™ sonicator (Diagenode). Cleared supernatants were further diluted with 2.2mL IP dilution buffer, precleared with rabbit IgG then incubated overnight at 4°C with 5–10 µg of antibody. Antibody-chromatin complexes were recovered using protein G-agarose beads (Roche). After washing, immunoprecipitated DNA was eluted from beads, crosslinks reversed, and DNA purified using phenol-chloroform-isoamyl alcohol. ChIP libraries were prepared by a commercial supplier (Novogene). Donor cells used in each experiment are listed in Table S1.

### ChIPmentation

ChIPmentation (CM) was carried out as described (Schmidl et al., 2015) with minor modifications. Biological triplicate experiments were performed for TFs except where noted otherwise. Five million nuclei were resuspended in 100μL sonication buffer (10mM Tris pH 8.0, 2mM EDTA, 0.25% SDS), sonicated for 10 cycles in a Bioruptor Pico™ sonicator (Diagenode) and diluted 1:1.5 ratio in equilibrium buffer (10mM Tris pH 8.0, 233mM NaCl, 1.66% Triton X-100, 0.166% sodium deoxycholate, 1mM EDTA). Cleared supernatants were incubated overnight at 4°C with 2µg of antibody, and antibody-chromatin complexes recovered using protein A/G magnetic beads. To improve signal to noise ratio for ERG CM only, we used a modified pull down protocol with two major differences: 1) nuclei lysates were not cleared by centrifugation after sonication, and 2) anti-ERG antibody was pre-conjugated to magnetic protein A/G beads, and then incubated with nuclear lysate overnight at 4°C to recover antibody-chromatin complexes.

After extensive washing, bead-bound complexes were resuspended in tagmentation mixture (25µL reaction containing 1 µL of enzyme in 1X buffer, Illumina) and incubated at 37°C for 25minutes. Crosslinking was reversed and DNA purified using a MinElute PCR Purification Kit (Qiagen). Barcoding/adapter primers (Table S7) and KAPA™ HiFi HotStart Ready Mix (Roche) were used to amplify libraries; the number of PCR cycles used was empirically determined for each reaction. Amplified DNA was purified then size selected using AMPure XP beads (Beckman) and sequenced using a standard Illumina 2 × 150bp PE pipeline (Novogene).

### HiC and HiChIP

HiC and HiChIP libraries were generated using the Arima Genomics HiC+ kit (Arima cat#A101020) respectively. Duplicate experiments were performed for HiC and HiChIP. Nuclei were lysed and chromatin digested with a restriction enzyme cocktail prior to end-filling with biotinylated nucleotides and ligation of proximal ends. For HiChIP, ligated fragments were then immunoprecipitated with the H3K27ac antibody. Biotinylated fragments were enriched and sheared prior to library preparation which was performed using Accel NGS 2S Plus DNA Library kit (Swift Biosciences).

### Bioinformatic processing

Analyses were run using default parameters for each tool unless otherwise indicated. Bigwig files were visualized using the UCSC browser (Kent et al., 2002). Reads were aligned to the GRCh38 genome accession GCF_000001405.26 (https://www.ncbi.nlm.nih.gov/assembly/GCF_000001405.26/). Heatmaps were plotted using seaborn unless otherwise indicated.

### Processing ATAC sequencing data

SRA files were downloaded from GEO (GSE74912), converted to fastq format, and aligned to the GRCh38 genome. Mitochondrial and duplicate reads were removed (respectively), reads shifted to account for Tn5-mediated excision, peaks called, and bigwig files generated (pipeline: SRA tools - BWA - samtools - picard MarkDuplicates - alignmentSieve - macs2 with a minimal threshold p-value of 1×10^-5^ - deeptools bamCoverage)(Li and Durbin, 2009; Li et al., 2009). To obtain a composite of all accessible regions in HSPCs, ATAC peaks from HSC, CMP, GMP, and MEP were merged using bedtools (Quinlan and Hall, 2010).

### Processing ChIP sequencing data

ChIP fastq files were analysed as single end data. fastq files were processed to remove adapter sequences (cutadapt)(Martin, 2011), trimmed to 100 bp (Trim Galore), then aligned to GRCh38 (BWA). Reads were sorted (samtools) and duplicates removed (picard MarkDuplicates), then bam files from replicates merged (samtools merge). IgG bam files from the four cell types were merged to generate a single IgG control. Peaks were called (macs2 with a minimal threshold p-value of 1 × 10^-5^ and using the IgG track as the control), and RPKM-normalized bigwig files generated (deeptools bamCoverage) (Ramirez et al., 2016). Composite plots showing ChIPseq signal at specific genomic regions were plotted in python as previously described (Thoms et al 2021).

### Motif enrichment analysis

Motif enrichment analysis was performed using the FIMO tool from the MEME analysis suite (Grant et al., 2011) using ETS, GATA, RUNX, and E-Box motifs sourced from JASPAR (Castro-Mondragon et al., 2022) as a position weight matrix.

### Analysis of combinatorial binding

Genomic locations with occupancy of multiple heptad TFs were identified by intersecting ChIP peak coordinates (bedtools intersect). To assess the significance of each combination we performed a bootstrapping analysis essentially as previously described (Beck et al 2013). Briefly, we derived an expected value for each combination using the merged ATAC peak set of 85,117 peaks, then compared the observed and the mean of the expected values, normalised to the expected standard deviation and calculated a z-score metric to denote significance of combinatorial events.

### Analysis of HiC and HiChIP data

HiChIP and HiC fastq files were processed and mapped to GRCh38, then PCR duplicates removed and contact matrices generated from the merged valid-pairs files (HiC-Pro hicpro2juicebox) (Servant et al., 2015). Contact matrices (.hic files) were visualized using juicebox (Durand et al., 2016). HOMER was used to identify compartments and TADs from balanced HiC-Pro contact matrices. The first principal component (PC1) was generated using runHiCpca.pl at 50 kb bins. In addition, H3K4me3 and H3K27me3 bed files from the respective cell types were used to assign accurate compartment labelling. To further identify TADs, findTADsAndLoops.pl function was used to generate TAD calls for each replicate separately – and merge2Dbed.pl to generate a union of TADs identified in each replicate. HiChIP contact matrices were used to generate interaction pairs at 5 kb resolution (MaxHiC)(Alinejad-Rokny et al., 2022) and the WashU browser was used for loop visualization (Li et al., 2019). Most interactions spanned distances >10 kb.

High confidence interactions (FDR ≤ 0.01) were used to generate a final list of promoter–regulator interactions. To map promoter-regulator interactions at heptad gene loci we identified HiChIP fragments which overlapped known promoters. Distal fragments that were linked to these promoters were intersected with ATAC peaks from the relevant populations to precisely map the contact region within 5kb HiCHIP fragments. Contact regions were named according to their linear genomic distance upstream (-) or downstream (+) from the transcriptional start site (TSS).

### Visualization of gene regulatory networks (GRNs)

GRNs were visualized with BioTapestry (Longabaugh, 2012) using ChIPseq peak calls and HiChIP-derived promoter–regulatory links to construct the network maps.

### Identifying differentially bound regions

Candidate regulatory elements (REs) were defined as regions displaying combinatorial binding of heptad factors with a positive z-score, indicating that the combination is observed at higher frequency than expected by random chance. DiffBind was used to identify regions showing a significantly higher (FDR ≤ 0.05) combinatorial TF signal in one cell type compared to all others (Stark and Brown, 2011). Two criteria were used for linking differentially bound REs with genes: the presence of a d-RE at a gene promoter, or of a HiChIP fragment that was in turn linked to a gene promoter.

### Analysis of bulk RNAseq data

Fastq files and count tables were downloaded from GEO (GSE75384) and fastq files aligned to GRCh38 (STAR) (Dobin et al., 2013). edgeR was used to normalise the count table and calculate log_2_ CPM values (Robinson et al., 2010) then derive a z-score of RNA expression.

### Analysis of single cell RNAseq data

SCANPY (Wolf et al., 2018) was used to process existing single-cell RNA sequencing data (Setty et al., 2019). The SCANPY score_genes tool was used to generate a score for our gene sets, which was ultimately plotted on the original tSNE map generated by those authors.

### TF occupancy at specific gene regulatory regions

Cell-type-specific gene lists were curated from the literature (Table S4). Promoter regions for these genes were defined as the ATAC peak occurring 1 kb upstream of the TSS, and distal fragments that were linked to these promoters in HiChIP datasets were intersected with ATAC peaks from the relevant populations to precisely map the contact region and define the distal regulatory element. Any promoters lacking a looped distal region were excluded from this analysis. To determine the heptad TF signal for each distal region, we added together the average signal across each region from log_2_-normalised bigwig tracks from each TF (bigWigAverageOverBed). If a looped distal regulator contained more than one ATAC peak, the TF signal from each peak region was averaged. We performed *k*-means clustering on the derived data using SciPy (Virtanen et al., 2020) and plotted the resulting heatmaps with seaborn.

### Clustering analysis

The merged set of ATAC peaks which represent open chromatin regions across HSPCs were annotated using ChIP (heptad, PU.1, CTCF, ATAC, H3K27ac, H3K4me3, and H3K27me3) and ATAC signal from each individual cell type to create a dataframe of 85,100 rows and 52 columns, and the regions clustered with SCANPY using the Louvain method (Blondel et al., 2008; Wolf et al., 2018). To identify regulatory regions preferentially used in specific cell types we compared TF signal at each region in HSC versus GMP, HSC versus MEP, and GMP versus MEP. We classified regions with log_2_ fold change > 2 for each heptad factor in a cell type as cell-specific-regions.

### Machine learning analysis to predict cell type

We trained models using the R package XGBoost (Chen and Guestrin, 2016). Briefly, we read a table of motif counts across the individual cell-specific-regions and took 70% of the peaks at random as the training set. We removed motifs with low variability and retained the remaining 30% of peaks as the test set. During the training, a series of decision trees were created such that a “loss function” was reduced (binary logistic in our case), to minimize cell type prediction error. Post training, prediction was performed on the test set. SHapley Additive exPlanation (SHAP) scores were calculated for every motif and peak used in the training set to indicate their respective contribution to the classification. A positive SHAP score for any given motif indicates that the presence of that motif in a region increases the probability that region belongs to the target cell type while a negative score indicates that the presence of that motif in a region increases the probability that region belongs to the background set (i.e., one of the other cell types). We then ranked motifs according to their importance by adding the absolute SHAP scores for every motif. To identify the direction of enrichment, we calculated the mean number of counts of every motif in the peaks that come from the target cell type or from the background peaks separately; if the mean was higher in the peaks from the target cell type those motifs were indicated as enriched in the target cell type and vice versa for the background set.

### Data availability

Raw and processed sequencing files have been uploaded to GEO with accession #XXXXXX.

