## Supplementary figures and legends for "Cell Type-Specific Regulation by a Heptad of Transcription Factors in Human Hematopoietic Stem and Progenitor Cells"

**Subramanian et al Supplemental Figures and Legends**

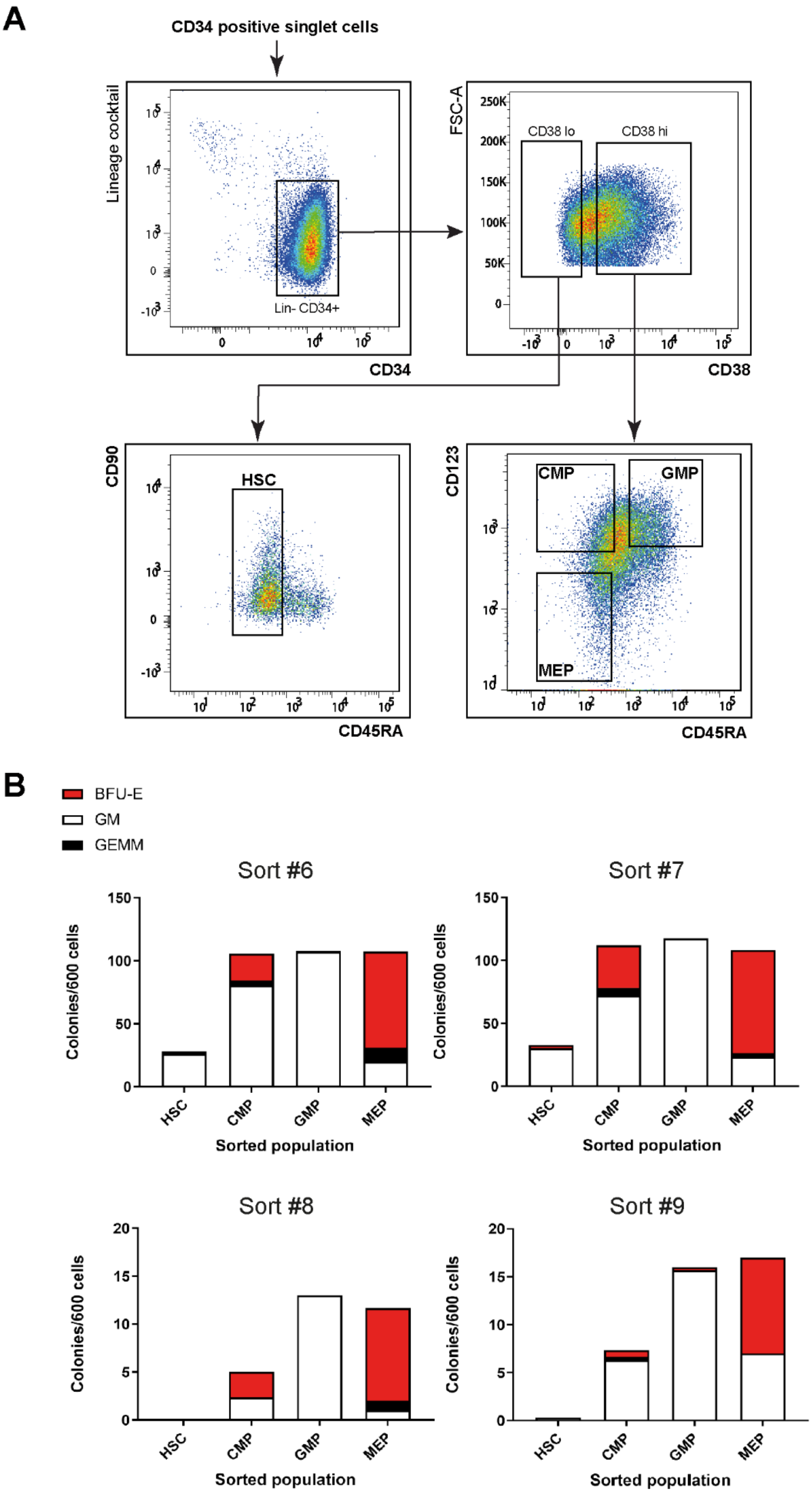
A
CD34 positive singlet cells
Lineage cocktail
Lin- CD34+
CD34
FSC-A
CD38 lo
CD38 hi
CD38
CD90
HSC
CD45RA
CD123
CMP
GMP
MEP
CD45RA
B
BFU-E
GM
GEMM
Sort #6
Sort #7
Sort #8
Sort #9
Colonies/600 cells
HSC
CMP
GMP
MEP
Sorted population

Figure S1

**Figure S1** ***Workflow for deriving HSPC fractions and verifying their identity***. **A)** Gating strategy and representative sort gates for isolation of HSC, CMP, GMP, and MEP. **B)** Representative CFU assays using sorted HSC, CMP, GMP, and MEP populations.

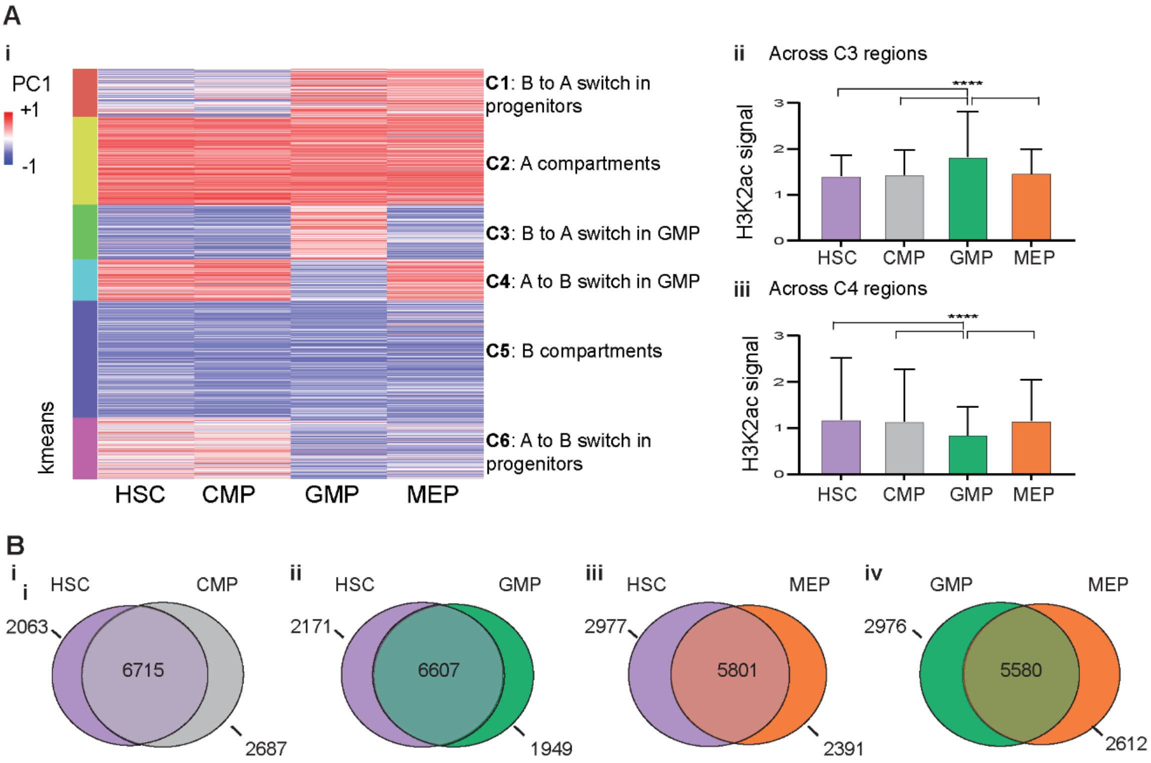


Figure S2

**Figure S2** ***Compartment switching and conserved TADs along blood stem cell differentiation***. **A)** i) A *k*-means clustered heatmap of the PC1 values showing compartmental switches taking place between the four HSPC populations. Average H3K27ac signal in ii) Cluster C3 regions, and in iii) Cluster C4 regions among the four cell types. Significance scores were calculated using pairwise t tests ($p < 0.0001$). **B)** Pairwise comparisons of topological domain boundaries between i) HSC and CMP, ii) HSC and GMP, iii) HSC and MEP, and iv) GMP and MEP (domains identified using HOMER).

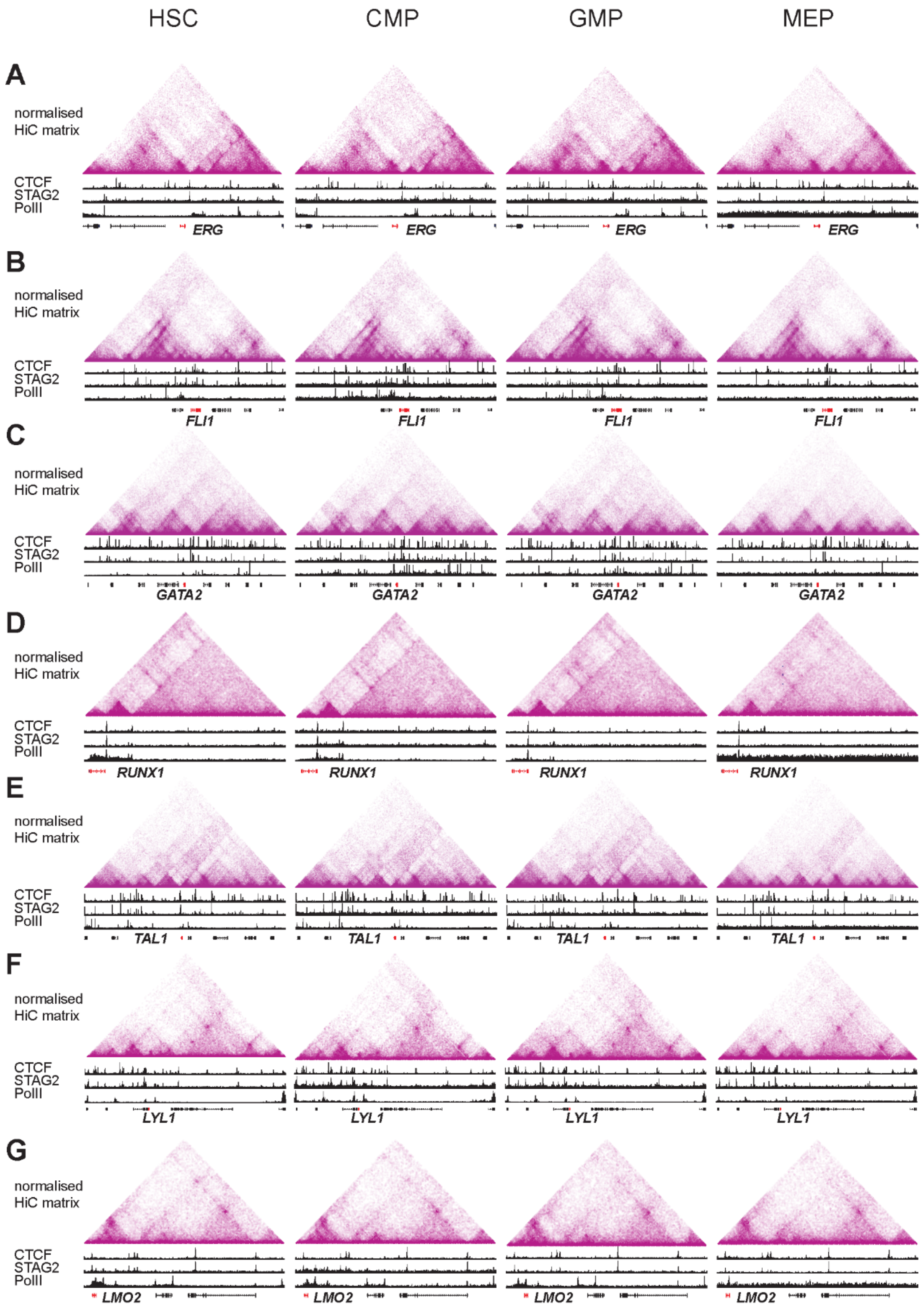
HSC
CMP
GMP
MEP
A
B
C
D
E
F
G
normalised HiC matrix
CTCF
STAG2
PolII
ERG
FLI1
GATA2
RUNX1
TAL1
LYL1
LMO2

Figure S3

**Figure S3.** ***The genome architecture at heptad gene regulatory loci is conserved across HSPC subsets.*** Normalised HiC contact matrices at 10 kb resolution located at individual heptad genes' regulatory loci – **A)** *FLI1* locus (GRCh38 chr11:128511084-128978507), **B)** *ERG* locus (GRCh38 chr21:37370238-39198738), **C)** *GATA2* locus (GRCh38 chr3:128262936-128761435), **D)** *RUNX1* locus (GRCh38 chr21:34758869-36011624), **E)** *TAL1* locus (GRCh38 chr1:47168881-47340728), **F)** *LYL1* locus (GRCh38 chr19:12787014-13852204), and **G)** *LMO2* locus (GRCh38 chr11:33831641-34445745), in HSC, CMP, GMP, and MEP respectively. Accompanying each triangular plot are ChIP-seq tracks showing CTCF-, STAG2-, and PolII-normalised signal.

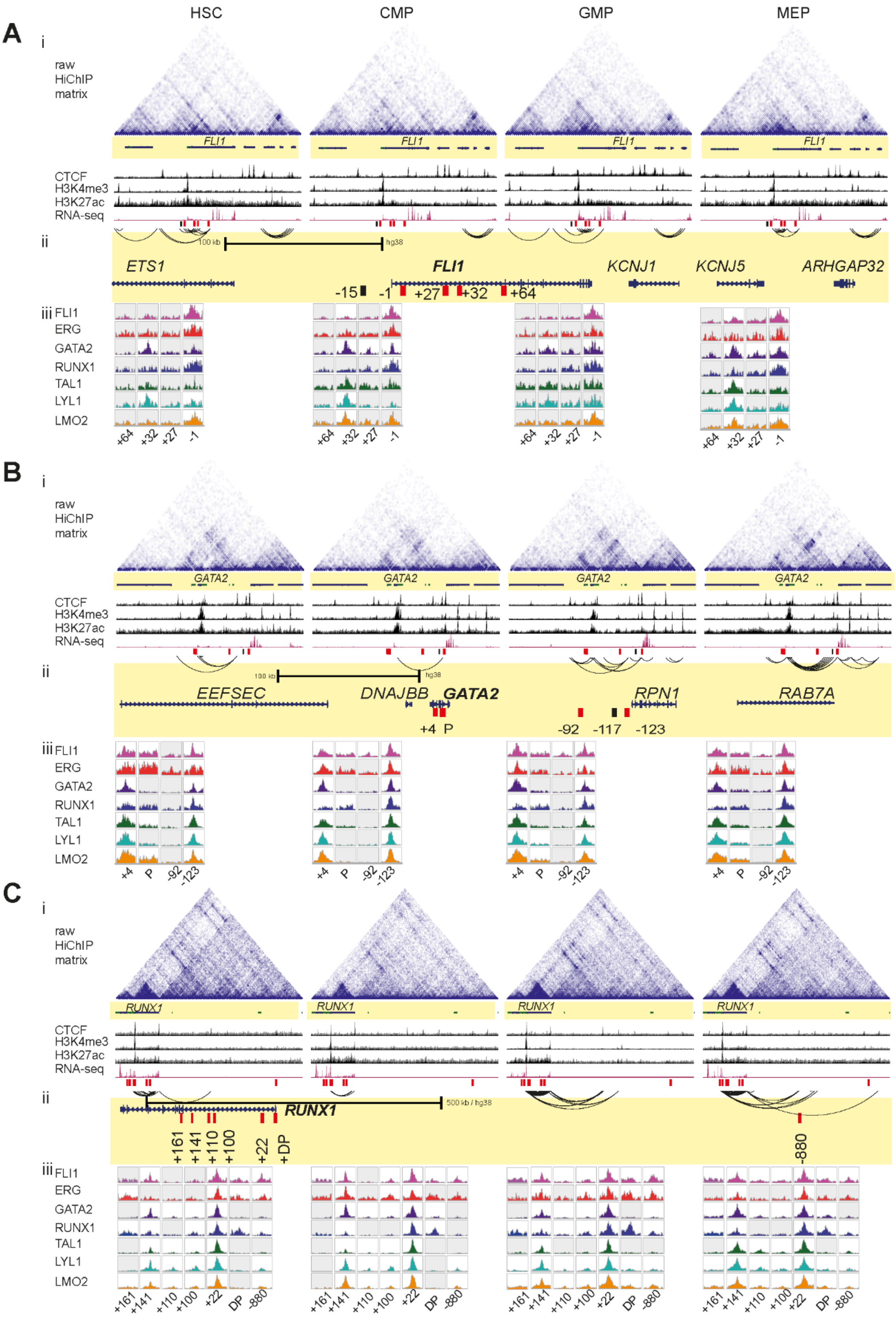

A
B
C
HSC
CMP
GMP
MEP
i
ii
iii
raw HiChIP matrix
CTCF
H3K4me3
H3K27ac
RNA-seq
FLI1
ERG
GATA2
RUNX1
TAL1
LYL1
LMO2
100 kb
hg38
ETS1
FLI1
KCNJ1
KCNJ5
ARHGAP32
-15
-1
+27
+32
+64
EEFSEC
DNAJBB
GATA2
RPN1
RAB7A
+4
P
-92
-117
-123
RUNX1
500 kb / hg38
+161
+141
+110
+100
+22
+DP
-880
DP


Figure S4

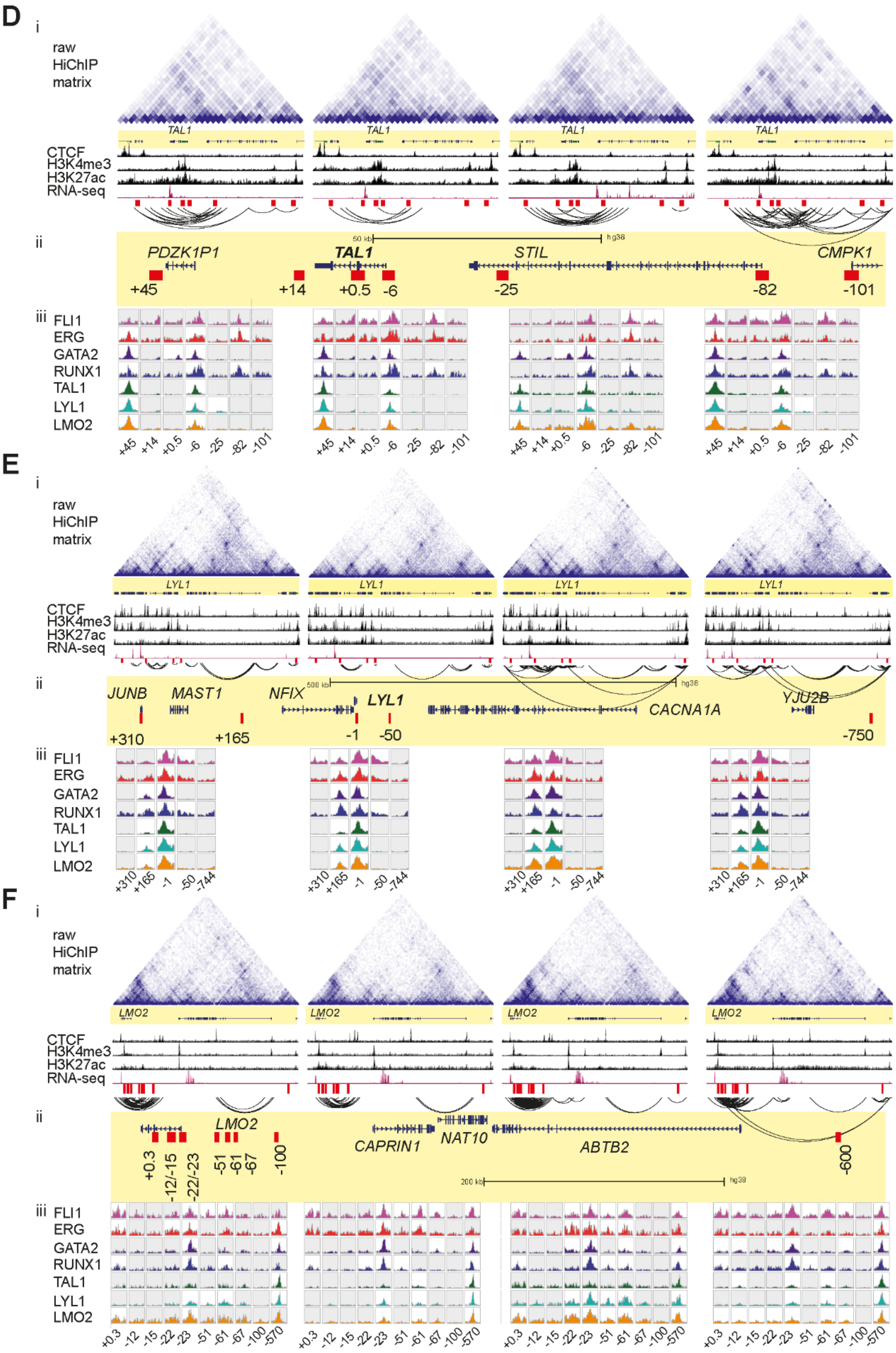
D
i
raw HiChIP matrix
TAL1
CTCF
H3K4me3
H3K27ac
RNA-seq
ii
PDZK1P1
TAL1
STIL
CMPK1
+45
+14
+0.5
-6
-25
-82
-101
iii
FLI1
ERG
GATA2
RUNX1
TAL1
LYL1
LMO2
E
i
raw HiChIP matrix
LYL1
CTCF
H3K4me3
H3K27ac
RNA-seq
ii
JUNB
MAST1
NFIX
LYL1
CACNA1A
YJU2B
+310
+165
-1
-50
-750
iii
FLI1
ERG
GATA2
RUNX1
TAL1
LYL1
LMO2
F
i
raw HiChIP matrix
LMO2
CTCF
H3K4me3
H3K27ac
RNA-seq
ii
LMO2
CAPRIN1
NAT10
ABTB2
+0.3
-12/-15
-22/-23
-51
-61
-67
-100
-600
iii
FLI1
ERG
GATA2
RUNX1
TAL1
LYL1
LMO2

Figure S4

**Figure S4.** ***H3K27ac HiChIP identifies cell-type specific interactions between heptad gene promoters and potential regulatory regions.*** **A)** i) Raw HiChIP contact matrix, CTCF, H3K4me3, H3K27ac, IgG, RNA-seq, and significant H3K27ac HiChIP interactions (FDR ≤ 0.01) at the *FLI1* locus (GRCh38 chr11:128511084-128978507). ii) Magnified view of the *FLI1* locus, with potential regulators looping to the promoter shown in red and potential regulators engaged in indirect regulatory activities shown in black. iii) FLI1, ERG, GATA2, RUNX1, TAL1, LYL1, and LMO2 peaks at the regulatory regions defined in ii, are shown. **B)** i) Raw HiChIP contact matrix, CTCF, H3K4me3, H3K27ac, IgG, RNA-seq, and significant H3K27ac HiChIP interactions (FDR ≤ 0.01) at the *GATA2* locus (chr3:128,262,936-128,761,435). ii) Magnified view of the *GATA2* locus, with potential regulators looping to the promoter shown in red and potential regulators engaged in indirect regulatory activities shown in black. iii) FLI1, ERG, GATA2, RUNX1, TAL1, LYL1, and LMO2 peaks at the regulatory regions defined in ii, are shown. **C)** i) Raw HiChIP contact matrix, CTCF, H3K4me3, H3K27ac, IgG, RNA-seq, and significant H3K27ac HiChIP interactions (FDR ≤ 0.01) at the *RUNX1* locus (GRCh38 chr21:34758869-36011624). ii) Magnified view of the *RUNX11* locus, with potential regulators looping to the promoter shown in red. iii) FLI1, ERG, GATA2, RUNX1, TAL1, LYL1, and LMO2 peaks at the regulatory regions defined in ii, are shown. **D)** i) Raw HiChIP contact matrix, CTCF, H3K4me3, H3K27ac, IgG, RNA-seq, and significant H3K27ac HiChIP interactions (FDR ≤ 0.01) at the *TAL11* locus (GRCh38 chr1:47,168,881-47,340,728). ii) Magnified view of the *TAL11* locus, with potential regulators looping to the promoter shown in red. iii) FLI1, ERG, GATA2, RUNX1, TAL1, LYL1, and LMO2 peaks at the regulatory regions defined in ii, are shown. **E)** i) Raw HiChIP contact matrix, CTCF, H3K4me3, H3K27ac, IgG, RNA-seq, and significant H3K27ac HiChIP interactions (FDR ≤ 0.01) at the *LYL1* locus (GRCh38 chr19:12787014-13852204). ii) Magnified view of the *LYL1* locus, with potential regulators looping to the promoter shown in red. iii) FLI1, ERG, GATA2,

RUNX1, TAL1, LYL1, and LMO2 peaks at the regulatory regions defined in ii, are shown. **F)** i) Raw HiChIP contact matrix, CTCF, H3K4me3, H3K27ac, IgG, RNA-seq, and significant H3K27ac HiChIP interactions (FDR $\leq$ 0.01) at the *LMO2* (GRCh38 chr11:33831641-34445745). ii) Magnified view of the *LMO2* locus, with potential regulators looping to the promoter shown in red. iii) FLI1, ERG, GATA2, RUNX1, TAL1, LYL1, and LMO2 peaks at the regulatory regions defined in ii, are shown. Only those HiChIP interactions where both interacting ends were found at the given locus are shown. In addition the ChIP-seq peaks shown are RPKM-normalised and white boxes indicate presence of a computationally called ChIP-seq peak at the specific region.

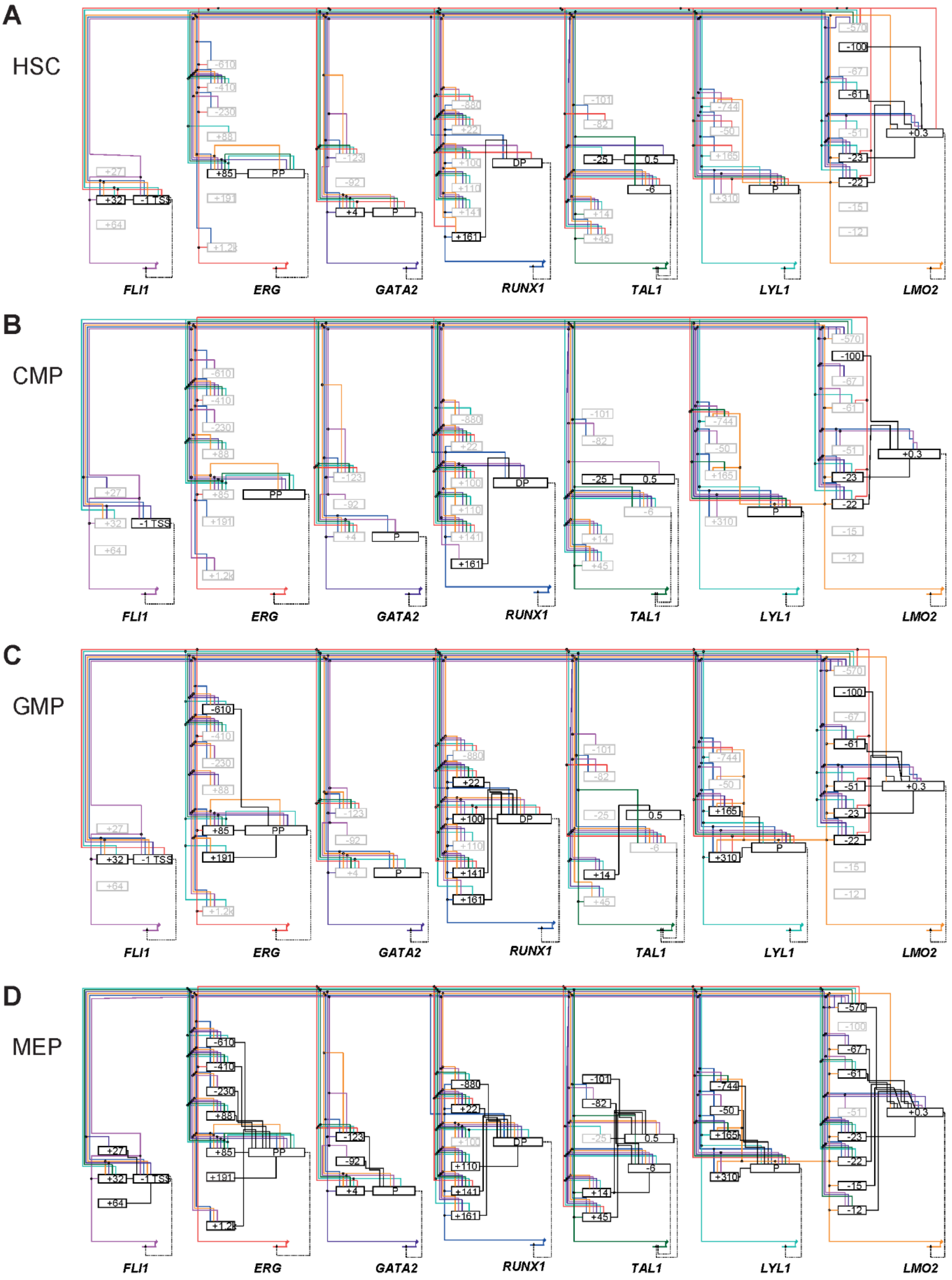


Figure S5

**Figure S5.** ***Gene regulatory network maps of the heptad genes in HSPC subsets.*** Heptad GRNs in **A)** HSC, **B)** CMP, **C)** GMP, and **D)** MEP, constructed using BioTapestry software. Boxes in bold show active regulators, and their interaction with respective promoters marked with solid black lines. Solid coloured lines indicate heptad factors binding to regulatory regions (FLI1-pink, ERG-red, GATA2-purple, RUNX1-dark blue, TAL1-green, LYL1-aqua, and LMO2-orange), while dashed lines link regulatory sub-circuits to individual genes.

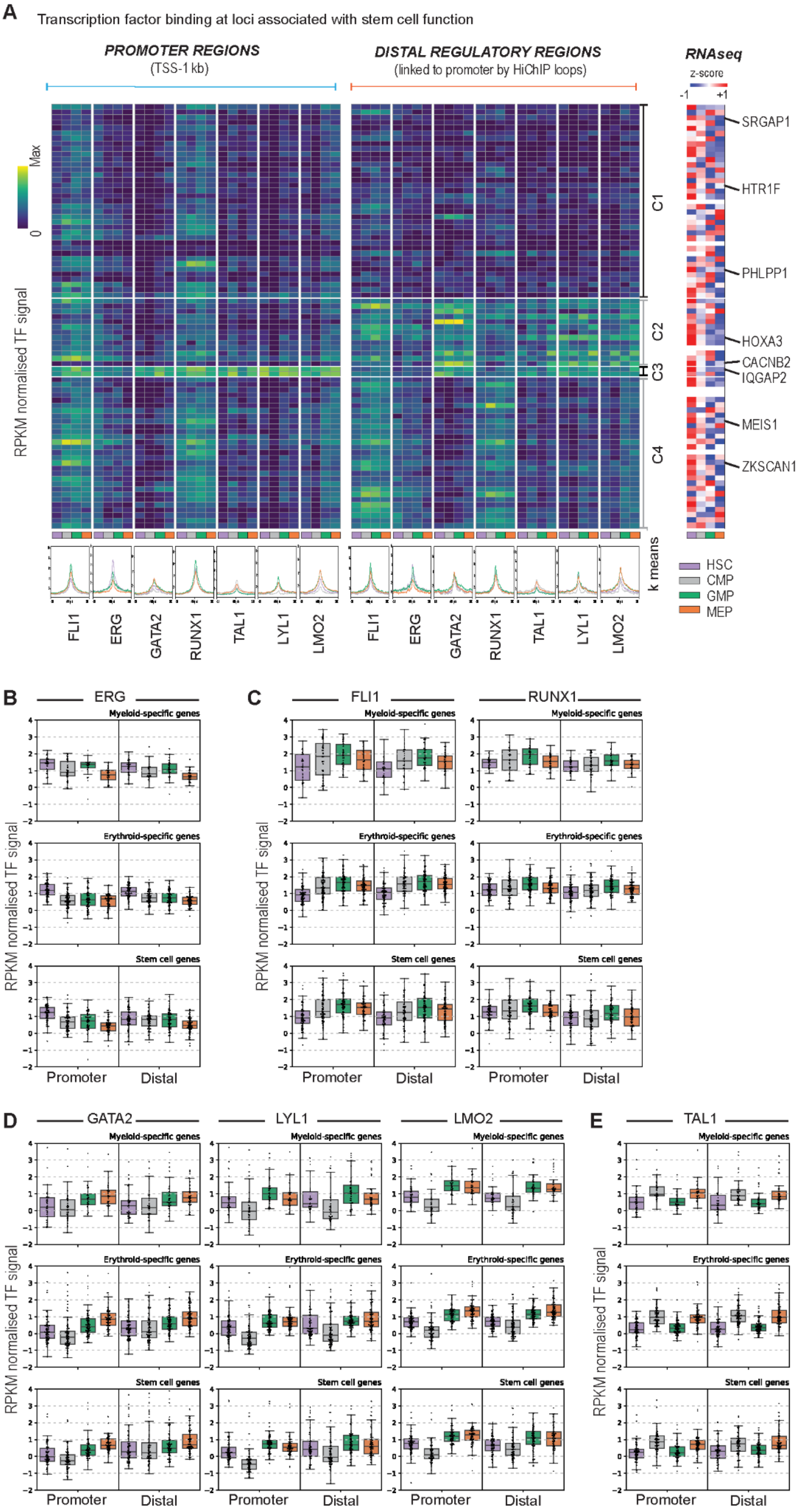

A
Transcription factor binding at loci associated with stem cell function
PROMOTER REGIONS
(TSS-1 kb)
DISTAL REGULATORY REGIONS
(linked to promoter by HiChIP loops)
RNAseq
z-score
RPKM normalised TF signal
Max
0
C1
C2
C3
C4
k means
SRGAP1
HTR1F
PHLPP1
HOXA3
CACNB2
IQGAP2
MEIS1
ZKSCAN1
HSC
CMP
GMP
MEP
FLI1
ERG
GATA2
RUNX1
TAL1
LYL1
LMO2
B
ERG
C
FLI1
RUNX1
D
GATA2
LYL1
LMO2
E
TAL1
Myeloid-specific genes
Erythroid-specific genes
Stem cell genes
Promoter
Distal


Figure S6

**Figure S6.** ***Cell-type specific binding patterns of heptad factors identified across stem cell specific genes*****. A)** Genes associated with stem cell function. *Left:* k-means clustered heatmaps of TF binding intensity at promoters and distal regulatory regions. Profile plots show normalised signal for each TF in each cell type at the regions depicted in the heatmap. *Right:* z-score normalised heatmaps of RNA-seq counts (GSE75384) for the corresponding gene in each cell type. **B-E)** Normalised TF signal at all promoters and distal regulatory regions for myeloid, erythroid, and stem cell genes. P-values for all pairwise comparisons (paired t-test) are shown in Table S4. **B)** Boxplots showing normalised ERG signal at promoters and distal regulatory regions of myeloid, erythroid, and stem cell genes. **C)** Boxplots showing normalised FLI1 and RUNX1 signal at promoters and distal regulatory regions of myeloid, erythroid, and stem cell genes. **D)** Boxplots showing normalised GATA2, LYL1, and LMO2 signal at promoters and distal regulatory regions of myeloid, erythroid, and stem cell genes. **E)** Boxplots showing normalised TAL1 signal at promoters and distal regulatory regions of myeloid, erythroid, and stem cell genes.

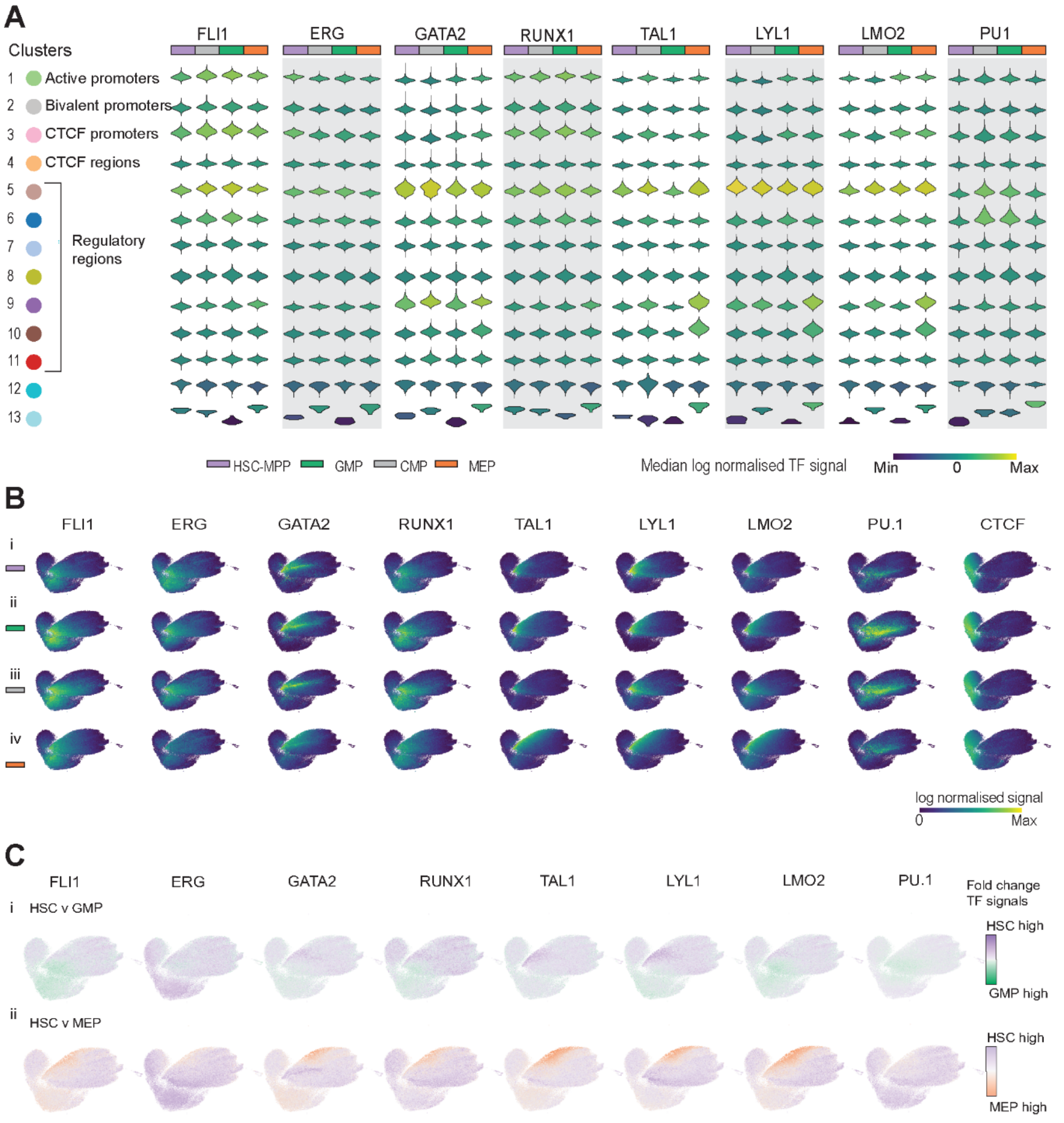


Figure S7

**Figure S7.** ***Transcription factor signal enrichment at ATAC regions in HSPCs.*** **A)** Individual violin plots show normalized transcription factor signals for FLI1, ERG, GATA2, RUNX1, TAL1, LYL1, and LMO2, in each of the 13 derived clusters for the four cell types studied. **B)** UMAPs showing heptad-factor-, PU.1-, and CTCF-normalized signals at accessible regions in i) HSC, ii) CMP, iii) GMP, and iv) MEP. **C)** UMAPs colored based on log2 fold change of binding of heptad transcription factors and PU.1 in pairwise comparisons between i) HSC and GMP, and ii) HSC and MEP.
